# A novel framework for inferring parameters of transmission from viral sequence data

**DOI:** 10.1101/302331

**Authors:** Casper K Lumby, Nuno R Nene, Christopher J R Illingworth

**Affiliations:** Department of Genetics, University of Cambridge, Cambridge, UK; Department of Applied Mathematics and Theoretical Physics, University of Cambridge, Cambridge, UK

## Abstract

Transmission between hosts is a critical part of the viral lifecycle. Recent studies of viral transmission have used genome sequence data to evaluate the number of particles transmitted between hosts, and the role of selection as it operates during the transmission process. However, the interpretation of sequence data describing transmission events is a challenging task. We here present a novel and comprehensive framework for using short-read sequence data to understand viral transmission events. Our model describes transmission as an event involving whole viruses, rather than independent alleles. We demonstrate how selection and noisy sequence data may each affect inferences of the population bottleneck, and identify circumstances in which selection for increased viral transmission may or may not be identified. Applying our model to data from a previous experimental transmission study, we show that our approach grants a more quantitative insight into viral transmission, inferring that between 2 to 6 viruses initiated infection, and allowing for a more informed interpretation of transmission events. While our model is here applied to influenza transmission, the framework we present is highly generalisable to other systems. Our work provides new opportunities for studying viral transmission.

## Introduction

Understanding viral transmission is a key task for viral epidemiology. The extent to which a virus is able to transmit between hosts determines whether it is likely to cause sporadic, local outbreaks, or spread to cause a global pandemic [1,2]. In a transmission event, the transmission bottleneck, which specifies the number of viral particles founding a new infection, influences the amount of genetic diversity that is retained upon transmission, with important consequences for the evolutionary dynamics of the virus [3,4].

Recent studies have used genome sequencing approaches to study transmission bottlenecks in influenza populations. In small animal studies, the use of neutral genetic markers has shown that the transmission bottleneck is dependent upon the route of transmission, whether by contact or aerosol transmission [5,6]. In natural human influenza populations, where modification of the virus is not possible, population genetic methods have been used to analyse bottleneck sizes. Analyses of transmission have employed different approaches, exploiting the observation or non-observation of variant alleles [7] or using changes in allele frequencies to characterise the bottleneck under a model of genetic drift [8–11]. A recent publication improved this latter model, incorporating the uncertainty imposed upon allele frequencies by the process of within-host growth [12]. Two studies of within-household influenza transmission have provided strikingly different outcomes in the number of viruses involved in transmission, with estimates of 1-2 [13] and 100-200 [14] respectively.

Another focus of research has been the role of selection during a transmission event; this is important in the context of the potential for new influenza strains to become transmissible between mammalian hosts [15,16]. Studies examining transmissibility have assessed the potential for different strains of influenza to achieve droplet transmission between ferrets under laboratory conditions [17–20]; ferrets provide a useful, if imperfect, model for transmission between humans [21,22]. The application of bioinformatic techniques to data from these experiments has identified ‘selective bottlenecks’ in the experimental evolution of these viruses [23,24], whereby some genetic variants appear to be more transmissible than others. In these studies, selection has been considered in terms of the population diversity statistic *π*; changes in *π*_*N*_/*π*_*S*_, the ratio between non-synonymous and synonymous diversity, have been used to evaluate patterns of selection across different viral segments.

We here note the need for a greater clarity of thinking in the analysis of viral transmission events. For example, analysis of genetic variants in viral populations shows that synonymous and non-synonymous mutations both have fitness consequences for viruses [25,26]; the use of synonymous variants as a neutral reference set may not hold. More fundamentally, in an event where the effective population size is small, the influences of selection and genetic drift may be of similar magnitude [27]. However selection is assessed, this implies a need to separate stochastic changes in a population from selection, especially where a transmission bottleneck may include only a small number of viruses [5,13,28]. The attribution of a change in diversity to the action of selection or the attribution of changes in allele frequencies to genetic drift could both potentially be flawed. Given the increasing availability of sequence data, more sophisticated tools for the analysis of viral transmission are required.

Here we note three challenges in the analysis of data from viral transmission events. Firstly, selection can produce changes in a population equivalent to those arising through a neutral population bottleneck [29] (Figure 1A), making it necessary to distinguish between the two scenarios. A broad literature has considered the simultaneous inference of the magnitude of selection acting upon a variant along with an effective population size [30–35]. However, such approaches rely on the observation of an allele frequency at more than two time points so as to distinguish a deterministic model of selection (with an implied infinite effective population size) from a combined model of selection with genetic drift; such approaches cannot be directly applied to the analysis of viral transmission.

**Figure 1.**
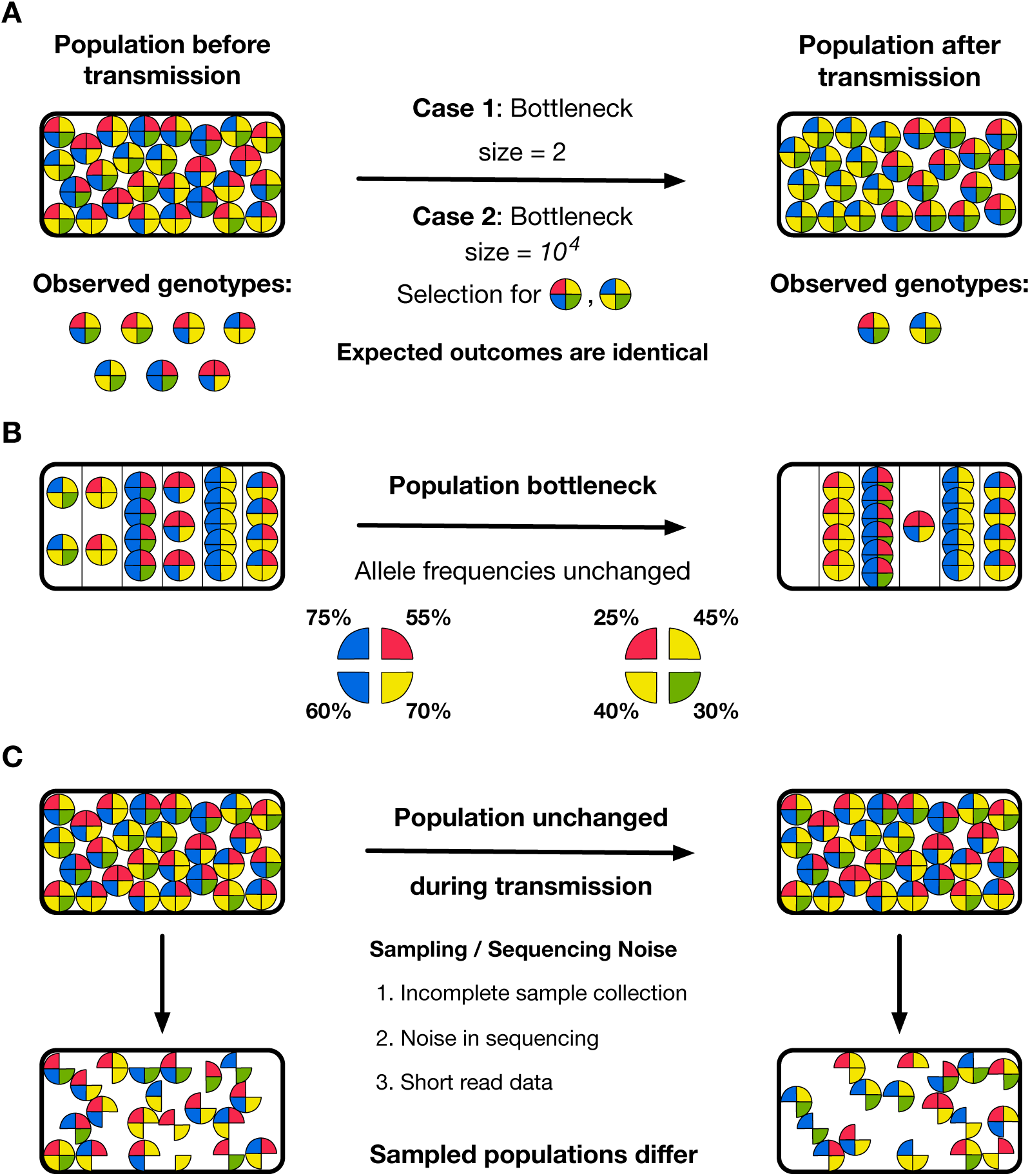
Challenges arising in the inference of transmission bottlenecks from viral sequence data. Circles represent idealised viral particles characterised by four distinct alleles. **A.** Reductions in population diversity cannot necessarily be attributed unambiguously to either a population bottleneck, or the action of selection. In the illustrated case, either a tight bottleneck without selection or a large bottleneck with strong selection could explain the change in the population during transmission. **B.** Straightforward statistics describing a population may generate misleading inferences of population bottleneck size. In the illustrated case, the genetic structure of a population is changed by a population bottleneck during transmission, but the frequency of each allele within the population does not change; an inference of bottleneck size derived from single-locus statistics would incorrectly be very large. **C.** Noise arising from the process of collecting and sequencing data is likely to produce differences between the observed populations, even in the event that the composition of the viral population was entirely unchanged during transmission.

Secondly, inferences of transmission events need to account for the haplotype structure of viral populations, whereby whole viruses, rather than sets of independent alleles, are transmitted (Figure 1B). The low rate of homologous recombination in segments of the influenza virus [36, 37] implies that viral evolution proceeds at the haplotype level [38]; competition occurs between collections of linked alleles, or segments, rather than the individual alleles themselves. Under such circumstances, fitter variants do not always increase in frequency within a population [39–41]. Calculations of genetic drift, which are often derived from the evolution of independent variants [42], need to be adjusted to account for this more complex dynamics.

Thirdly, noise in the measurement of a population may influence the inferred size of a transmission bottleneck (Figure 1C). A broad range of studies have examined the effect of noise in variant calling and genome sequence analysis [43–50]; more recently formulae have been proposed to measure the precision with which allele frequencies can be defined given samples from a population [51–53]. Where small changes in allele frequencies are used to assess a population bottleneck, it is important to separate the effects of noise in the measurement of populations from genuine changes in a population.

We here describe a novel method for the inference of population bottlenecks which resolves the above issues. Our approach correctly evaluates changes in a population even where the data describing that change is affected by noise. It explicitly accounts for the haplotype structure of a population, utilising the data present within short sequence reads. Further, where these factors can be discriminated, our method distinguishes between the influences on the population of selection and the transmission bottleneck. Studies of viral evolution have highlighted the potential for payoffs between within-host viral growth and transmissibility [54]; given sufficient data we can evaluate how selection operates upon each of these two phenotypes. Our model extends previous population genetic work on bottleneck inference to provide a generalised model for the analysis of data spanning viral transmission events.

## Results

### Model outline

In the recent literature, the term ‘bottleneck’ has been applied to describe a reduction in the genetic diversity of a population (e.g. [55]), whether arising from selection or a numerical reduction in the size of a population. Here, we define a ‘bottleneck’ more strictly as a neutral process whereby a finite number of viral particles from one population found a subsequent generation of the population, either within the same host, or across a transmission event from host to recipient. Selection then constitutes a modification to this process whereby some viruses, because of their genotype, have a higher or lower probability of making it through the bottleneck to found the next generation.

We applied a population genetic method to make a joint inference of the bottleneck size and the extent of selection acting during a transmission event. We consider a scenario in which a viral population is transmitted from one host to another, with samples being collected before and after the transmission event (Figure 2A). In our model viruses are categorised as haplotypes, that is, according to the variants they contain at a subset of positions in their genome. As such, the viral population is represented as a vector of haplotype frequencies; the population before transmission is represented by the vector ***q**^B^*. During transmission, a random sample of *N^T^* viruses are passed on to the second host to give the founder population ***q**^F^*. Selection for transmissibility, whereby genetic variants cause some viruses to be more transmissible than others, is described by the function *S^T^*. The potentially small size of the founder population means that the population evolves within the host under the influence of genetic drift to create the large post-transmission population ***q**^A^;* this process is approximated in our model by a population bottleneck of effective size *N^G^*. Selection acting for within-host growth may further alter the genetic composition of the population; this effect is described by the function *S^G^*. Our method thus allows for a discrimination to be made between selection for increased within-host replication and for increased viral transmissibility. Observations of the population are collected before and after transmission via a noisy sequencing process to give the datasets ***x**^A^* and ***x**^B^*. The extent of noise in the sampling and sequencing is characterised by the parameter *C*; a value of *C* = 0 would indicate that samples contain no information, while a large *C* corresponds to noiseless sampling, as in a binomial model.

**Figure 2.**
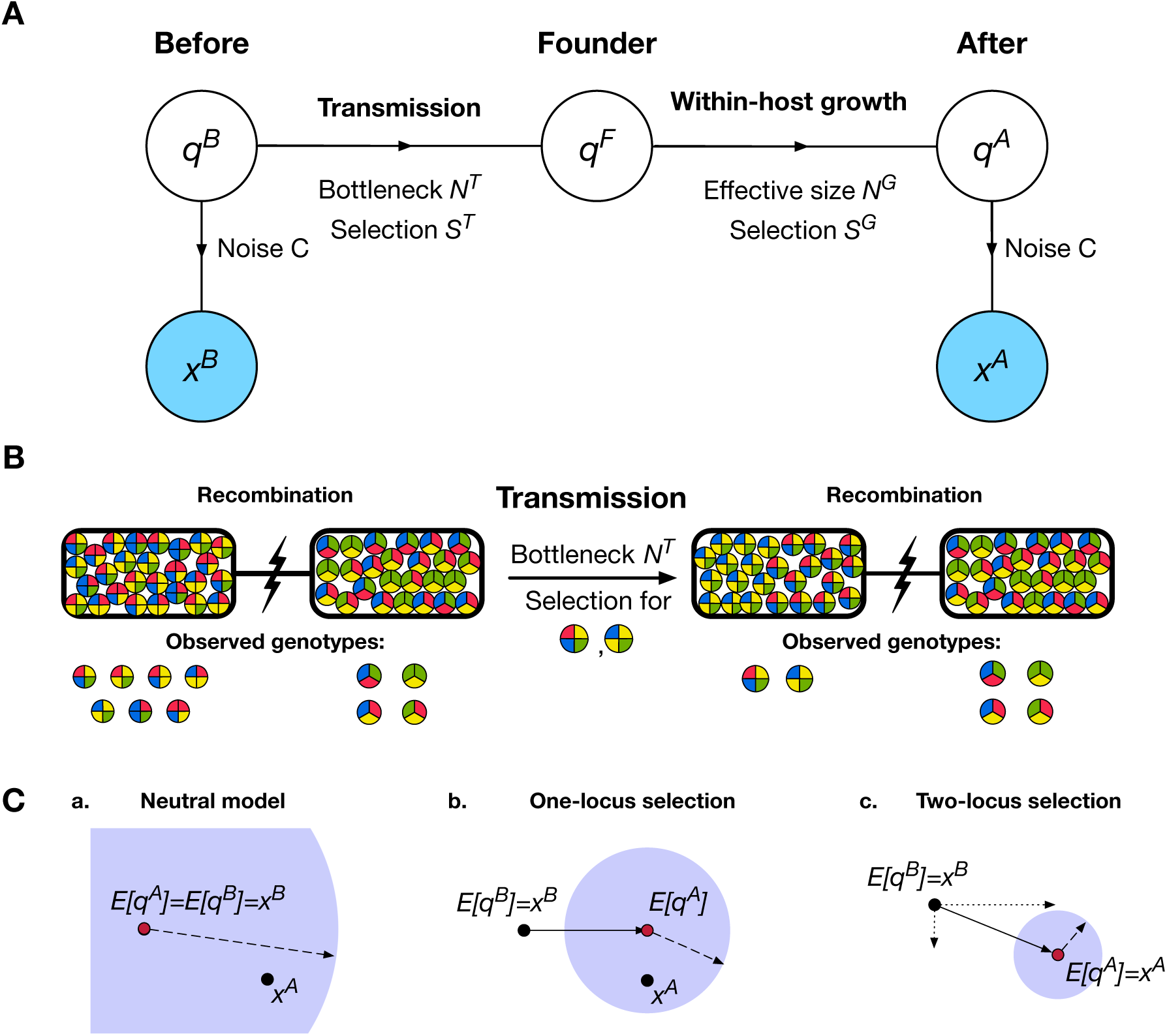
**A.** Basic model of transmission. A set of haplotypes exists at frequencies ***q**^B^* from which a noisy observation ***x**^B^* is made. During a transmission event, a total of *N^T^* viruses are transferred under the influence of selection *S^T^*, establishing an infection in the next host described by ***q**^F^*. Growth of the viral population within the host then occurs to produce the population ***q**^A^*, influenced by genetic drift (characterised by the effective population size *N^G^*) and selection *S^G^*. Sampling of the final population gives the second observation ***x**^A^*. **B.** Regions of the genome which are separated by recombination or reassortment are used to distinguish the effects of selection and a population bottleneck. Here, genetic diversity is reduced in one region but not in another; the preservation of diversity in the second region attributes this change to the action of selection on the first, rather than a shared population bottleneck. **C.** Models of neutrality and selection are compared, as illustrated in this simplified diagram. Black dots represent observations ***x**^B^* and ***x**^A^* while the red dot indicates the inferred expected position of ***q**^A^*. The solid line joining these (b,c) indicates the inferred action of selection, with dotted lines showing components of this vector (c). The blue circle represents the optimised variance in the position of ***q**^A^*; the length of it’s radius, shown as a dashed line, is inversely related to the bottleneck size. In the neutral case, the difference between observations is explained by the bottleneck alone. More complex models of selection fit ***q**^A^* more closely to ***x**^A^* and with reduced variance, giving higher inferred values of *N^T^*.

To summarise our approach, we note that both the transmission and within-host growth events can be represented as sampling processes, which may each be biased by the effect of selection. As such, given an estimate of the noise inherent to the sequencing process, and externally-derived estimates for *N^G^* and *S^G^*, we can calculate an approximate likelihood for the parameters *N^T^* and *S^T^* given the observations ***x**^B^* and ***x**^A^*. Maximising this likelihood gives an estimate for the size of the transmission bottleneck and the extent to which specific genetic variants within the pre-transmission population confer increased transmissibility upon viruses.

In our model we discriminate between changes in a population arising from selection and those arising due to the population bottleneck. This is achieved by considering regions of the genome between which recombination or reassortment has removed linkage disequilibrium between alleles (Figure 2B). As transmission involves whole viruses, the bottleneck *N^T^* is preserved between regions. Meanwhile, in the absence of epistasis, selection acting upon one region of the virus does not influence the composition of the population in other parts of the genome. As such, a cross-region calculation estimates both *N^T^* and the influence of selection. A model selection process [56] is used to distinguish models of neutral transmission from evolution under selection (Figure 2C). A full exposition of the model is given in the Methods section.

Here we used simulated data to evaluate the performance of our model under different circumstances. Having established the effect of sequencing noise on the inference of population bottlenecks, we demonstrate the ability of our method to correctly infer population bottlenecks from sequence data in the presence or absence of selection, and its ability to correctly identify variants conferring a benefit for viral transmissibility. We then applied our model to evaluate selection and population bottlenecks in a recently published viral transmission experiment [24]; our approach provides more precise inference of population bottlenecks in this case and discriminates between the influence of selection for within-host growth and viral transmission.

### Application to simulated data

#### Sequencing noise limits the maximum inferrable bottleneck

Application of our model to simulated data describing neutral population bottlenecks showed that a lack of sequencing noise is critical for the correct inference of large population bottlenecks (Figure 3). Noise in our study was considered in terms of the precision with which the frequency of a variant can be specified by viral sequence data. Our noise parameter *C* can be related to an ‘effective depth’ of sequencing [53], involving the absolute number of reads of a genome position, which specifies the depth of noise-free reads that would produce the same degree of certainty in an allele frequency. This statistic is dominated by the smaller of *C* and the absolute read depth; for large *C* this statistic is close to the absolute read depth, while as the number of reads describing a variant frequency becomes large, the effective depth tends to *C* + l.

**Figure 3.**
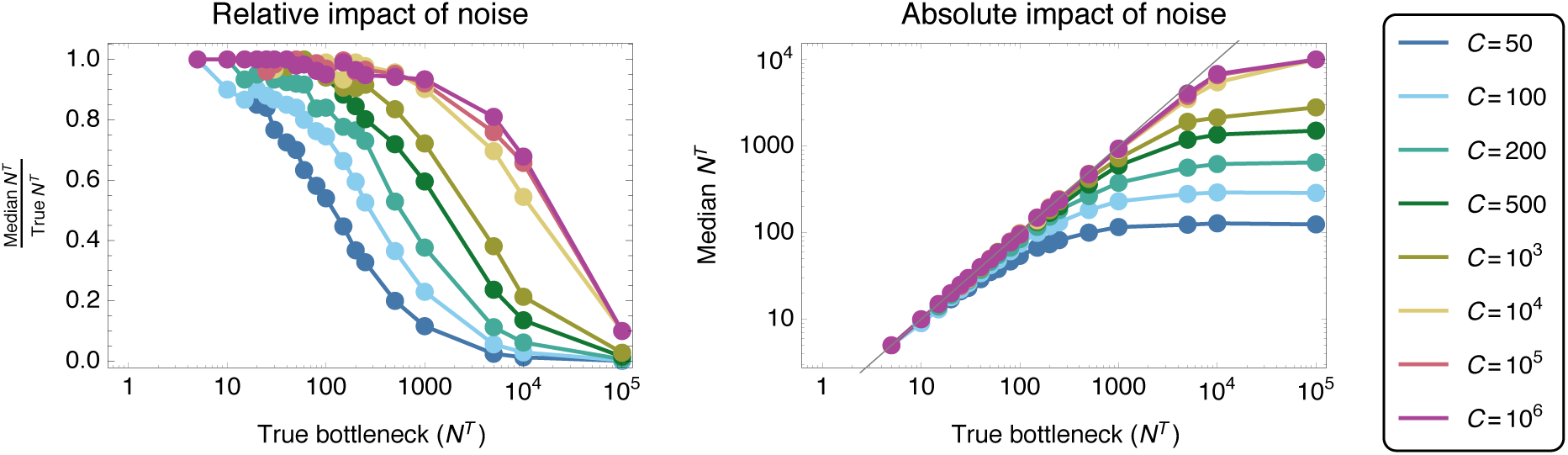
Influence of sequencing noise upon the ability to infer a population bottleneck size from genome sequence data. Median inferred bottlenecks are shown, calculated on the basis of 200 replicate simulations for each point. In the left-hand plot, a value of 1 indicates a correct bottleneck inference; in the right-hand plot, the absolute inferred bottleneck size is shown. Simulations were conducted under the assumption of selective neutrality, with no attempt to infer selection from the data.

Inferences of bottleneck sizes showed a limit on the inferred bottleneck size governed by noise in sequencing; where there was little noise in the data (i.e. at high values of C), a correct inference of the true population bottleneck was generally made. However, as noise increases, the inferred bottleneck reaches a plateau above which increases in the true bottleneck no longer affect the inferred bottleneck size. This result can be understood in terms of the extent to which the population bottleneck and noise contribute to the change in the viral population; where large numbers of viruses are transmitted, most of this signal is likely to result from noise. Here we note failures in the inferred bottleneck size even with very high *C*; these occur due to the finite read depth in our simulations, which averaged around 12,000. In these calculations a neutral method, in which selection was assumed to have no effect on the population, was used to make inferences from neutral simulations. A consistent value of *C* was used for simulation and inference purposes.

In a real dataset, the extent of noise may be unknown. Further investigation showed bottleneck estimation to be relatively robust to an incorrect estimate of the extent of noise in a dataset, except where the extent of noise was substantially overestimated (Supplementary Figure S1). In general, an underestimate of the extent of noise in a dataset led to an inferred bottleneck size that was marginally lower than the correct value, while an overestimate of the extent of noise led to an overestimate of the size of the bottleneck. Severe overestimation of noise led to dramatically incorrect inferences; as such, while noise limits the intrinsic potential of a method to identify large bottleneck sizes, underestimating the extent of noise is generally the more safer approach.

#### Variance in inferred transmission bottlenecks

Results from individual simulations showed that our method could discriminate between bottleneck sizes that differ by a factor of four or above (Figure 4). Obtaining precision in an estimated bottleneck or effective population size is inherently a difficult task, relying on the estimate of the extent of a stochastic effect from limited data [32]. Across 200 simulations, the interquartile range in an inferred bottleneck spanned close to 28% of the true bottleneck size, with inferred values spanning a range of approximately 140% of the correct bottleneck size. A slight underestimate in the bottleneck size for the case *N^T^* = 100 was consistent with the extent of noise in sequencing; here and in all subsequent simulations a value of *C* = 200 was used, representing an extent of noise that is readily achievable from short read sequencing [51,53]. In our inferences, while gross differences in bottleneck size can be identified, a high level of precision is difficult to obtain from sequence data alone.

**Figure 4.**
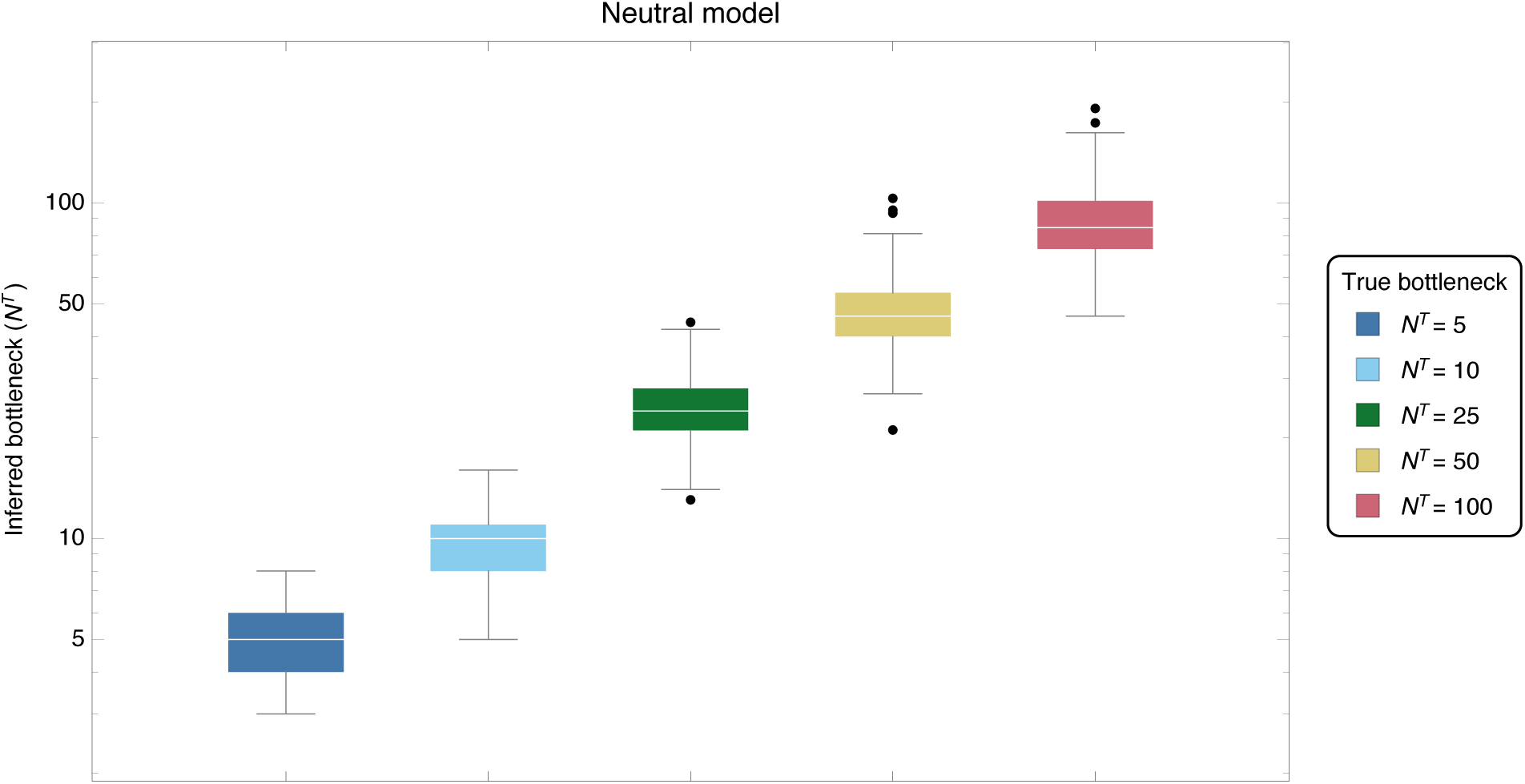
Inferred bottleneck sizes (*N^T^*) for true bottlenecks *N^T^* = {5,10, 25, 50,100}. Results were generated by applying a neutral inference model to neutral simulated data. Inferences are shown for 200 simulations at each bottleneck size.

#### Inference of population bottleneck sizes under selection for transmissibility

Inferences of bottleneck size showed a systematic underestimate of the bottleneck when selection affected a transmission event, but a method neglecting selection was used in the inference procedure (Figure 5). Simulations were conducted in which an allele at the third of five polymorphic loci in the HA segment of a simulated influenza virus increased the transmissibility of the virus according to a selection coefficient *σ*; this model of selection was applied for all subsequent simulations. In our simulations a value of *σ* = 1 is equivalent to a change in the frequency of a variant from 50% to 73% in a single transmission event. The relatively strong magnitudes of selection considered reflect the short period of time (a single generation) over which selection for increased transmissibility can act and the relatively small number of viruses likely to be involved in a transmission event.

**Figure 5.**
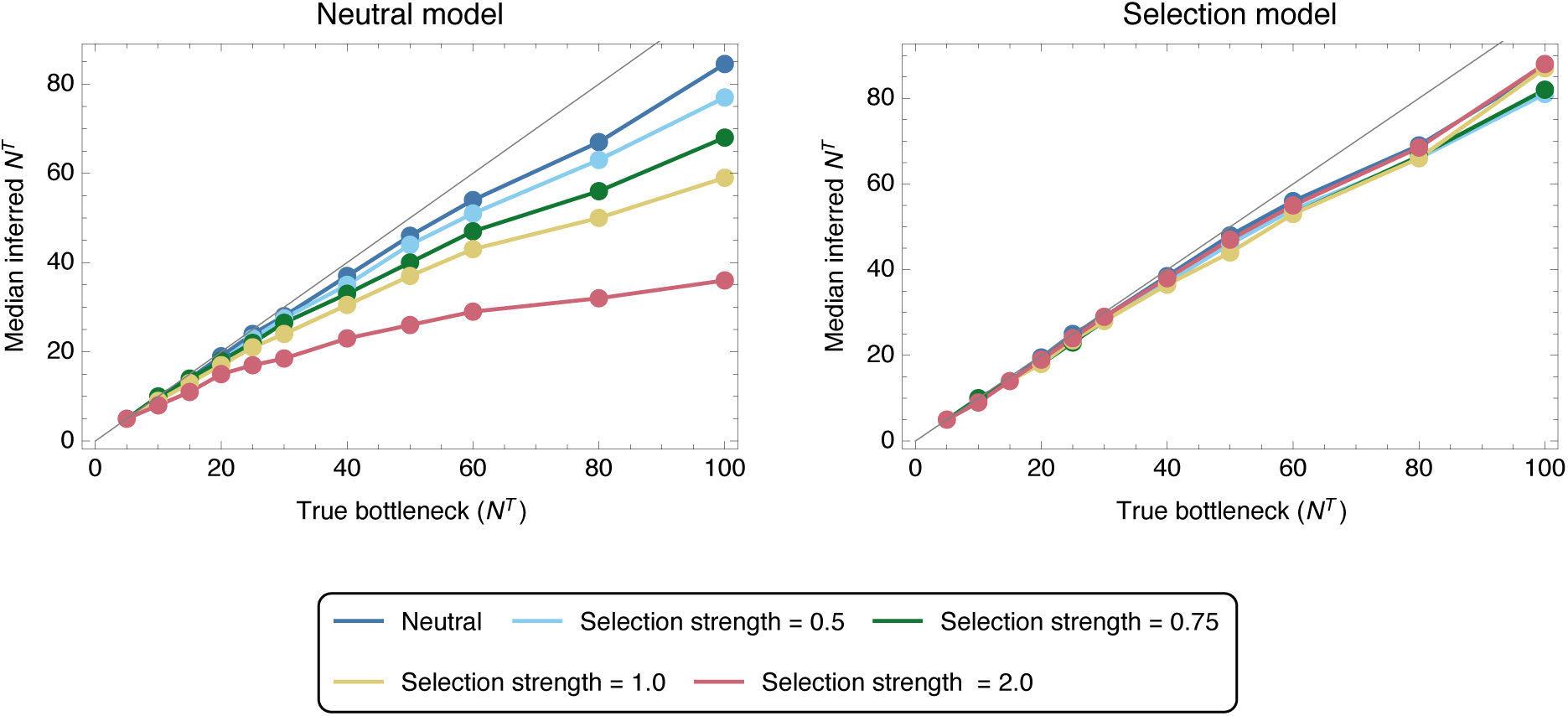
Median inferred bottleneck size from data simulating transmission with a single locus under selection of magnitude *σ* ∈ {0,0.5, 0.75,1.0, 2.0}. Inferences were made using either a neutral inference model, in which the effect of selection was assumed to be zero, or a model incorporating selection, which allowed the presence of selection to be inferred. Median inferences are shown from 200 simulations for each data point.

Inferences of population bottleneck were conducted using a neutral inference method, and with a model in which selection was not constrained to be zero. In the first case, ignoring selection led to an underestimation of the true bottleneck size by an amount which increased according to the magnitude of selection for transmissibility. Selection during transmission produces a shift in the expected composition of the viral population; if this shift is interpreted as occurring due solely due to a finite bottleneck a tighter bottleneck, inducing a larger stochastic change in the population, is inferred. This understanding explains the more pronounced underestimates achieved at larger bottleneck sizes; larger bottlenecks produce smaller stochastic changes in the population relative to the change induced by selection. When the full version of our model was run, allowing for a consideration of selection effects, the median bottleneck inferred from data under selection resembled that inferred from neutral data; the small shortfalls in the inference from neutral data are here explained by the influence of noise.

Repeated calculations performed for data describing multiple replicate transmission events gave similar inferred transmission bottlenecks. In each case sets of three replicate transmission events were simulated, transmitted populations sharing a common set of polymorphic loci in each segment. Median inferred values are shown in Supplementary Figure S2. Full results describing the range of inferred bottleneck sizes from both one- and three-replicate populations are shown in Supplementary Figures S3 to S6.

#### Identification of variants under selection

In contrast to measures of diversity, which attempt to associate selection with a gene or segment of a virus, our method was able to correctly identify specific variants conferring increased transmissibility. Success was more often achieved in cases for which selection was relatively strong and the transmission bottleneck was relatively large (Figure 6). Our process for distinguishing selection from neutrality (Figure 2C) can be tuned to identify a greater number of true variants under selection at the cost of making a greater number of false positive calls; here a conservative approach to identifying selection was applied. Under this approach we retained a false positive rate (inference of selection at an unselected locus) of 5% or less across the systems tested. At lower magnitudes of selection (*σ* ≤ 0.5), correctly identifying sites under selection was very difficult, though as selection became stronger (*σ* ≥ 1) loci under selection could be identified with greater accuracy. Where selection existed the ease with which it could be identified increased with increasing bottleneck size. Our results can again be understood with respect to the dynamics of the system. The bottleneck has a stochastic effect on the population of a magnitude inversely related to the number of viruses transmitted. Inferring the presence of selection requires the identification of changes in the population going beyond what would be expected under neutrality, biasing the population in the direction of the selected allele or alleles. However, stochastic effects can by chance distort the population in one direction or another by more than the expectation; this leads to false inferences of selection. Genuine changes resulting from selection become easier to identify when the changes are themselves larger (stronger selection) or where the magnitude of the stochastic effect is reduced (higher *N^T^*). In contrast to the inference of bottleneck size, data from replicate simulations led to a more dramatic change in the results, with the false positive rate falling to zero for bottlenecks with *N^T^* ≥ 15. The power of replicate experiments arises from the lower probability that stochastic effects will impose a consistent pattern of change upon multiple populations. While a larger-than-expected stochastic change in the frequency of a variant may occur in one system, leading to a false positive inference of selection, it is unlikely that the same pattern would recur across multiple replicates. The use of replicate experiments is therefore very powerful for identifying variants affecting transmissibility; while, under our conservative approach, not all variants under selection were identified, variants identified from replica data as being under selection were almost universally true positive calls.

**Figure 6.**
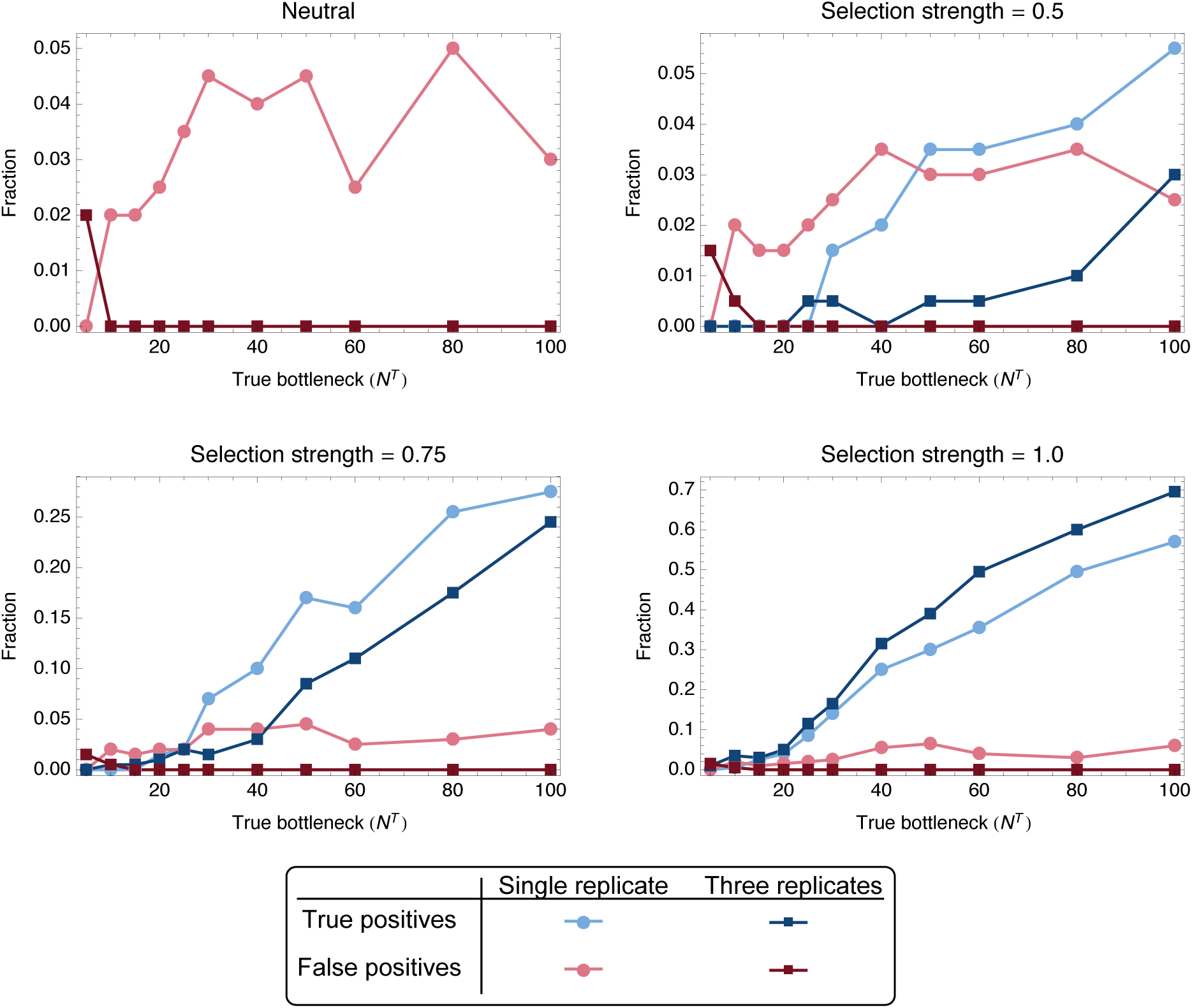
True and false positive rates of selection inference from 200 simulations of transmission events from single- and three-replicate systems with selective pressures of *σ* ∈ {0,0.5, 0.75.1.0}. True positives were defined as inferences for which selection was inferred for the selected locus in a system; false positives were defined as inferences for which selection was inferred at any neutral locus or for multiple neutral loci in the system.

#### Estimating the magnitude of a selected variant

Given the correct identification of selection acting for a specific variant, the inferred magnitude of selection was marginally overestimated, with an increased overestimate at smaller values of the transmission bottleneck *N^T^* (Figure 7). The mixture of deterministic and stochastic changes in the population explains this phenomenon; the population after transmission is equal to its expected value plus some stochastic change. In the event that the stochastic change is aligned with the direction of selection, the presence of selection is more likely to be inferred, while the additional change in that direction will give an over-estimate of selection. Conversely, if the stochastic change is in a direction opposed to the influence of selection, the presence of selection is less likely to be inferred. Thus, selection was disproportionately inferred to exist when stochastic changes in the population led to an overestimate of its magnitude. Inferences conducted on sets of replicate transmission events produced more accurate and more precise estimates of selection. For example given a bottleneck of *N^T^* = 100 and a true strength of selection of 0.75, the mean inferred selection from a single replicate was 0.98 with variance 0.048, while the mean inferred selection from three replicates was 0.87 with variance 0.013. (Supplementary Figure S7)

**Figure 7.**
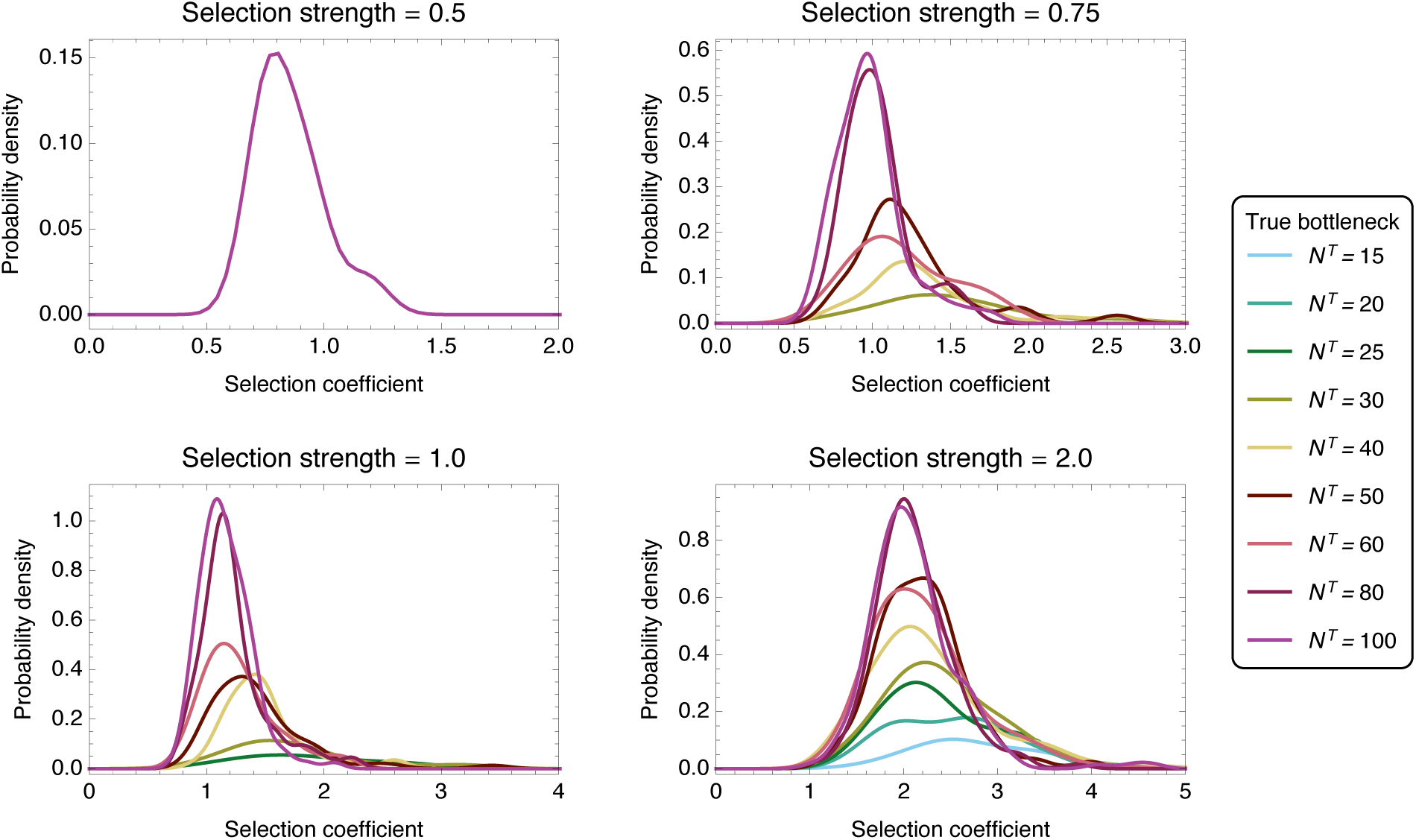
Probability distributions of inferred selection coefficients from 200 simulations of transmission events with selective pressures *σ* ∈ {0.5, 0.75,1.0, 2.0}. Distributions were constructed for bottleneck values where the inference of selection resulted in a true positive rate for identifying selected variants of above 5 %. Smooth kernel distributions were computed using a Gaussian kernel function defined on (0,10) and Silverman’s rule of thumb [57, p. 48] for the bandwidth size. Distributions were scaled such that their integral across the kernel range equalled the true positive rate.

### Application to an experimental dataset

We applied our approach to an influenza transmission dataset obtained by Watanabe et al. [58] and subsequently analysed by Moncla et al. [24]. This dataset provides high-resolution, whole-genome sequence data describing both the within-host evolution, and airborne transmission, of a 1918-like influenza virus, that became transmissible upon introduction of three key mutations, PB2 E627K, HA E190D and G225D.

This three-mutant strain was denoted ‘HA190D225D’ and successfully transmitted in one of three ferret transmission pairs. Isolation and subsequent growth in MDCK cells of viruses from the contact ferret of the successful transmission led to the generation of the ‘Mut’ strain, which transmitted in two of three instances. A previous analysis of these data using linked variants on the HA segment identified an increase in the diversity of the viral population during within-host growth, and respectively ‘loose’ and ‘stringent’ bottlenecks in the transmission of the two strains. In the transmission of the Mut strain, the fixation of sequence variants, potentially due to selection, was observed, while the observation of two out of three, rather than one out of three, successful transmissions suggested that the Mut virus may have evolved increased fitness for infection. Within and between hosts, segment-wide and localised measures of synonymous and non-synonymous sequence diversity *π* were used to assess the presence or absence of selection, leading to the conclusion that selection affected the system during transmission of the ‘Mut’ strain.

In our study, data from serial samples from the within-host populations were used to infer a fitness landscape for within-host growth for each of the two populations. Using a previously published approach [51] we inferred the presence of non-neutral change in the population in seven out of eight segments in the combined HA190D225D population, and in four out of eight segments in the combined Mut population. The inference of positive selection acting for multiple non-consensus viral haplotypes in the HA segment (Figure 8) explains the increase in sequence diversity previously observed. Further results are shown in Supplementary Figures S8 and S9 and in Supplementary Table S1.

**Figure 8.**
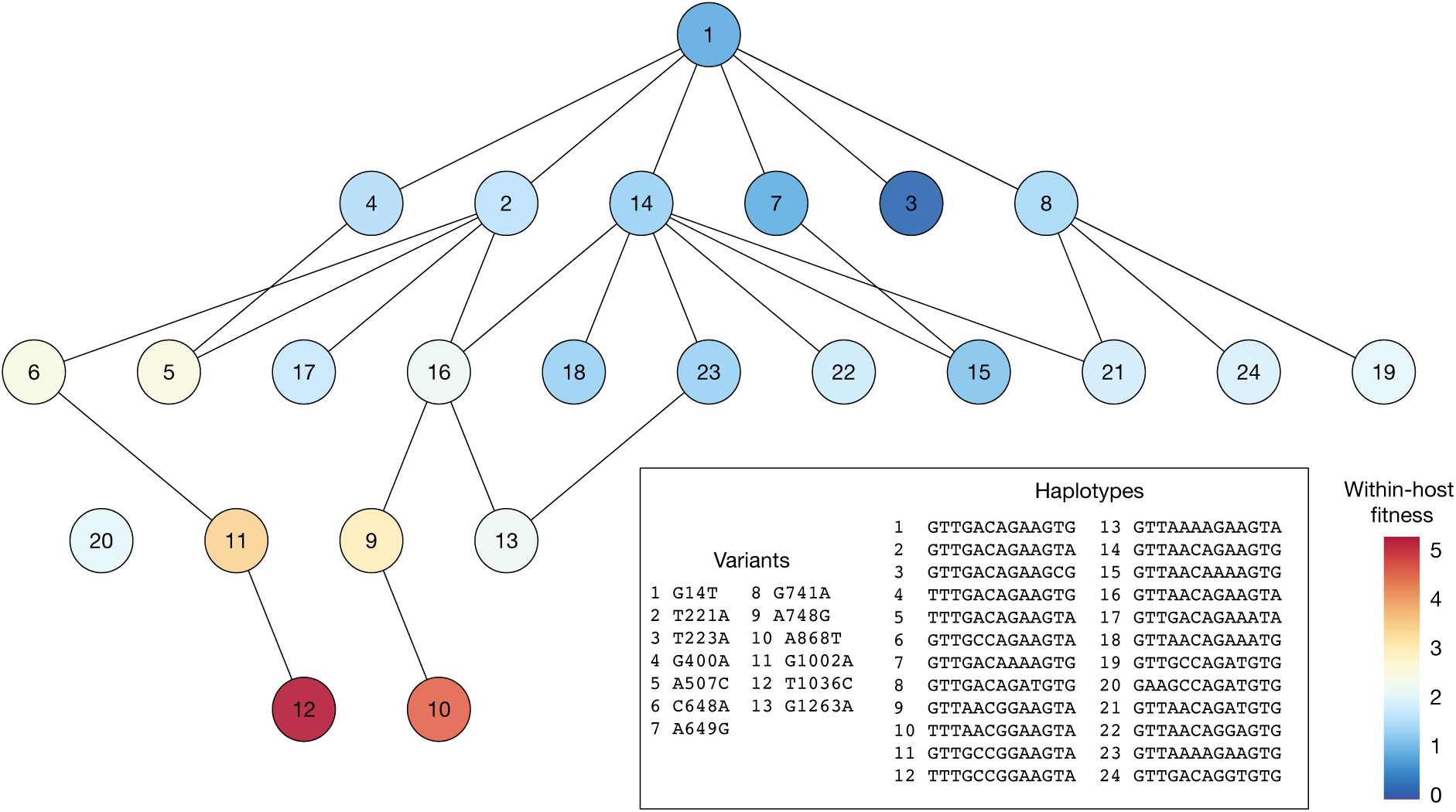
Inferred fitness landscape for within-host growth using data from the HA190D225D dataset. Viral haplotypes for which the inferred frequency rose above 1% in at least one animal are shown. Lines show haplotypes separated by a single mutation.

Applying our inference framework to the data identified narrow transmission bottlenecks in each case (Figure 9). In each of our calculations a set of statistical replicate inferences was produced, corresponding to different potential reconstructions of the population ***q**^B^* from the sequence data (see Methods). Within the HA190D225D population, our estimated bottlenecks ranged from 4 to 6, with a median bottleneck size of 5, while for the Mut calculations, our bottlenecks ranged from 2 to 37 and 2 to 8, with medians of 6 and 2 respectively. As such, no clear evidence was found that the HA190D225D transmission involved a greater number of particles than the Mut transmissions. Given the inclusion of the inferred within-host selection *S^G^*, no evidence was found for the existence of variants making the virus more or less transmissible, with selection being inferred in only a small number of the replicate calculations (Supplementary Figure S10). Increasing the frequency cutoff at which variants were included in the calculation led to small decreases in the inferred bottleneck sizes (Supplementary Figure S11).

**Figure 9.**
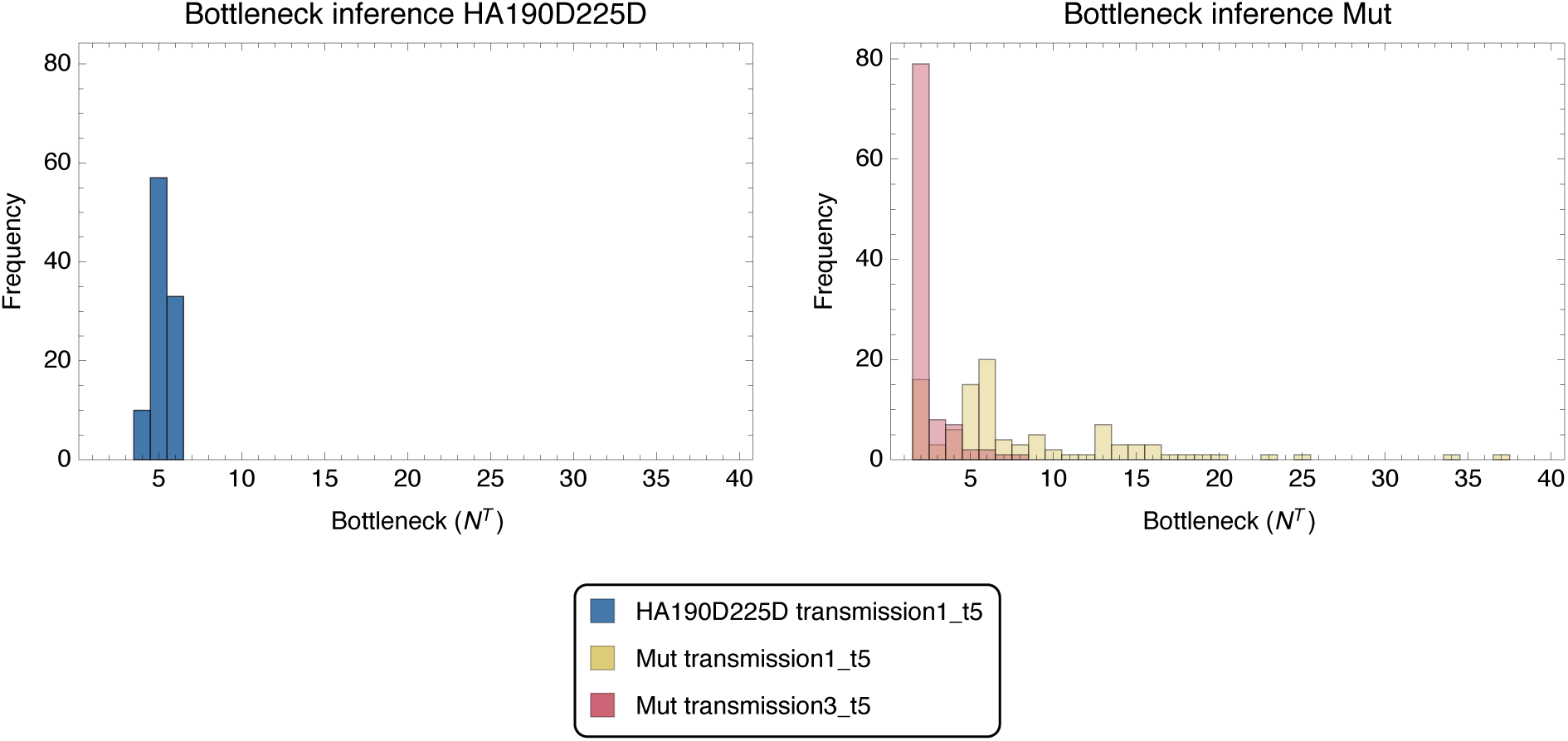
Histograms of bottleneck inferences for HA190D225D and Mut transmission pairs from 100 analysis seeds. A replicate inference method was employed for the Mut transmission pairs such that a common fitness landscape was imposed. The Mut transmission pairs may take different bottleneck values and have been plotted as an overlapping histogram.

## Discussion

We have here presented an approach for jointly inferring a population bottleneck size and selection for differential transmissibility from viral sequence data describing a transmission event. While basic sampling approaches to bottleneck inference have been improved by an accounting for drift during within-host viral growth [7,12–14], our approach additionally accounts for noise in genome sequence data, exploits partial haplotype data available from short-read sequencing, and separates the influence of a finite bottleneck from that induced by selection for increased transmissibility. In multiple studies, the transmission bottleneck has been found to be narrow during natural viral spread between hosts [59]. While we do not explicitly question these results, we note that both selection and noise in the measurement of sequence data can decrease the inferred bottleneck where sequence data is used for inference. Our approach is suitable for the analysis of acute infectious diseases such as influenza on the basis of a small number of observed transmission events; we note that where more substantial diversity is present in a within-host viral population, or where data is available from a large number of hosts in an outbreak, phylogenetic methods of evolutionary inference become of increasing value [60–62].

Applied to the analysis of data from a recent evolutionary experiment, our approach provides a greater precision in the inference of evolutionary statistics, leading to an alternative explanation for the data observed. Where data have previously been interpreted as implying differential transmission bottlenecks between strains, our approach infers bottlenecks of similar sizes. Furthermore, where evidence has been interpreted to suggest a differing extent of transmissibility between strains, our approach attributes changes in allele frequencies either to stochastic effects or to selection for increased host adaptation. Our result does not definitively prove the absence of differential transmissibility among the viruses involved in this study, but implies that data which might suggest differential transmissibility can be more parsimoniously explained in other ways.

Our study shows that the identification of variants conferring increased viral transmissibility is difficult when the number of transmitted viral particles is small. While improvements to our method may be achievable, this difficulty is fundamentally rooted in the nature of a transmission event; where a low number of virions transmit, the influence of stochastic processes become large, with variants fixing during transmission in a manner that cannot be distinguished from a selective sweep. The potential to infer the presence of selection increases at larger population sizes. The size of the transmission bottleneck in natural influenza populations is currently a subject of debate [13,14]; where experiments are conducted to assess viral transmissibility, steps taken to increase the number of particles transmitted would increase the power to infer differential transmissibility. We note that, unlike more general inferences of selection from changes in viral diversity, our approach evaluates selection in terms of specific variants conveying an advantage or disadvantage for transmission. Where broad measures of diversity are calculated across segments of a genome, the background of genetic diversity across a large number of positions may be hard to separate from changes at individual positions under the action of selection.

In the light of our study, we propose that the term used in some analyses of viral transmission, of a ‘selective bottleneck’ is ambiguous, failing on the one hand to distinguish changes in a population arising from selection and those occurring through stochastic change in the population, and on the other to distinguish between selection for more rapid within-host selection or for inherent viral transmissibility. While selection may act differently for these latter two phenotypes [54], their respective influences are intrinsically hard to separate from data. In this case, the completeness of the collected data, covering both within-host adaptation and between-host transmission, was necessary to evaluate the cause of evolutionary change.

Our framework may reduce the need for animals in viral transmission studies. One approach to exploring transmissibility (in influenza virus) has been the comparison, for different viruses, of the proportion of distinct animal pairs between which transmission occurs [63]. The statistical significance achievable in these studies is limited by the number of animal pairs that can be examined [64–66]. Furthermore, the comparison between one genotype and another may be confounded by viral heterogeneity, whereby each population contains a cloud of genetic diversity [23,67]. In our approach, while data from replicate transmission events is of value, it is the number of particles transmitted, rather than the number of replicates, that primarily informs inferences of transmissibility. Transmission of more viruses between fewer animals may provide a more efficient use of animal stocks.

In some situations, neutral markers or molecular barcodes may be added to a viral population in order to characterise bottleneck sizes [5,29]. While our method does not require the presence of such markers, its adaptation to include marker data would likely be straightforward, including in a calculation a further probabilistic term constraining the bottleneck size. Inference of selection for transmissibility could then be conducted under this constraint; the combination of whole-genome sequence data with such information could prove powerful for the study of viral transmission.

While we have here considered the transmission of influenza virus, very few steps of our approach would need to be altered for the method to be applied to another viral population. As detailed in the Methods section, it is only in accounting for genetic drift in the within-host growth of the virus that we make approximations relying on biological knowledge of the influenza virus; an alternative accounting for within-host expansion could be used. A second key assumption in the inference of selection is the existence of regions of the virus separated from each other by recombination or reassortment. This assumption would be preserved in some other viruses, as noted in observations of within-host HIV evolution [68], if not for all influenza populations [69]. Where a viral genome did not exhibit recombination, and only a single transmission event was observed, the neutral version of our method could be applied; in so far as we utilise partial haplotype data, and account for sequencing noise, our method would still provide an advantage over alternative methods.

Viral transmission is a critical component of disease and a key factor in viral evolution. In outlining a novel framework for the interpretation of data from viral transmission events we hope to bring a greater clarity to the population genetic theory of how these events operate and a greater power in the interpretation of experimental data, so as to engender a greater understanding of this important topic of research.

## Methods

### Notation and qualitative overview

We describe the viral population as a set of haplotypes, with associated frequencies, that changes in time during a transmission event. Given a number of (possibly non-consecutive) loci of interest in the viral genome, the set of haplotypes ***h*** = {*h_i_*} decribes a set of sequences having specific nucleotides at these loci. Within a viral population of finite size, the number of viruses with each haplotype *h_i_* is described by the vector ***n*** = {*n_i_*}. Frequencies of each haplotype within the population are denoted by the vector ***q*** = {*q_i_*}, while observations of the population collected via sequencing are denoted by the vector ***x*** = {*x_i_*}, where *x_i_* is the number of sampled viruses with haplotype *h_i_*.

The transmission event is now described according to the framework outlined in Figure 2. A population of viruses ***q**^B^* undergoes transmission with some bottleneck *N^T^*, creating a founder population with haplotype frequencies ***q**^F^* in the recipient. Selection influencing this transmission process is described by the function *S^T^*(***q***), which changes the frequency of haplotypes according to the relative propensity of each haplotype to transmit. Within the host, the viral population grows rapidly in number to create the population ***q**^A^*. During this growth process, genetic drift affects the population in a manner according to the effective population size *N^G^*. Observations of the system are made via genome sequencing of samples collected before and after transmission, and are denoted ***x**^B^* and ***x**^A^* respectively; the total numbers of sequence reads in each are denoted *N^B^* and *N^A^*. Given the observations ***x**^B^* and ***x**^A^*, we wish to estimate the size of the population bottleneck *N^T^* and the extent of selection for transmissibility *S^T^*.

During the process of growth between ***q**^F^* and ***q**^A^*, the population may be influenced by selection for within-host growth; this acts independently of selection for transmissibility [70], and is described by the function *S^G^*(***q***), which changes the frequencies of haplotypes according to their relative within-host growth rates. Selection for within-host growth is challenging to separate from selection for transmissibility; we here estimate this parameter independently from the transmission event itself.

### Likelihood framework

As the observations ***x**^B^* and ***x**^A^* are conditionally independent given ***q**^B^*, the joint probability of the system may be written as a product of individual probabilities

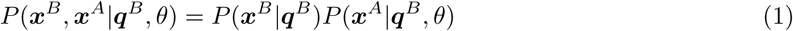

where *θ* represents the remaining variables in the system upon which only ***x**^A^* depends.

As an approximation to this likelihood, we split the inference into two calculations, first calculating a maximum likelihood for ***q**^B^* given ***x**^B^*, then inferring the transmission event from ***x**^A^* given ***q**^B^*. Noting the potential uncertainty in the inference of ***q**^B^*, we introduce a variance component so that ***q**^B^* may be regarded as a random variable rather than a fixed quantity. The process of breaking up the inference process greatly reduces the computational time required for our approach, without considerable cost to the accuracy of the results. Splitting the likelihood in this manner, and marginalising over unknown quantities, the likelihood can be written generically as

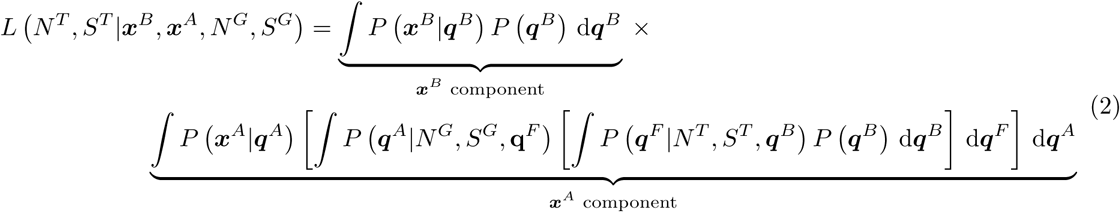

The first component of this likelihood, corresponding to the initial observation of the system, ***x**^B^*, represents a straightforward sampling of the system, drawing from a collection of viral haplotypes. Such a process can be modelled using a multinomial distribution. However, as is well known [53], next-generation sequence data is error-prone, such that less information is contained within the sample than would be contained in a multinomial sample of equivalent depth to the sample. A Dirichlet multinomial distribution may be used to capture this reduction of information, such that

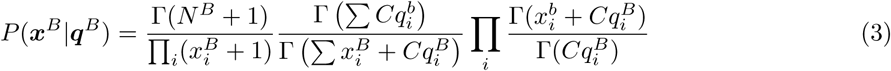

where *C*, which alters the variance of the distribution, characterises the extent of noise in the data. The parameter *C* can be estimated given independent observations of identical parameters, such as haplotype or single allele frequencies; in the application to experimental data, time-resolved variant frequencies derived from the sequence data were used for this purpose [51]

Considering the second component of the likelihood, the expression *P*(***x**^A^*|***q**^A^*) may be calculated in the same manner as in Equation 3 dependent upon the haplotype frequencies ***q**^A^*. The remaining parts of this component can also be described as sampling events. A sample of the population in the donor animal transmits to the recipient, generating a founder population. The founder population multiplies within the host to generate the final population ***q**^A^*. The ***x**^A^* component thus represents a compound of multiple sampling events. We will go on to describe the calculation of both components of the likelihood function. However, we first need to consider how selection is incorporated into our model.

#### Excursus: Modelling selection

Within our model, the functions describing selection are potentially complex, each having a number of parameters equal to the number of haplotypes in the system. In common with previous approaches to studying within-host influenza evolution [71] we adopt a hierarchical model of selection whereby the fitness of a haplotype is calculated from a set of one- or multi-locus components, describing the advantage or disadvantage of a specific nucleotide, or nucleotides, at a single locus or set of loci. Model selection is then used to identify the most parsimonious explanation of the data.

Formally, we denote the *j^th^* component of the haplotype *h_i_* as *h_ij_*, with *h_ij_* ∈ {*A*, *C*, *G*, *T*}. In a fitness model, a parameter is defined as the pair of values (*s_k_*, *g_k_*), where *s_k_* is a real number, denoting the difference in fitnesses of individuals with and without the allele [72], and *g_k_* is a vector of components *g_kj_* ∈ {*A*, *C*, *G*, *T*, –} denoting the haplotypes to which this selection applies. We now define

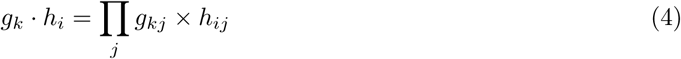

where

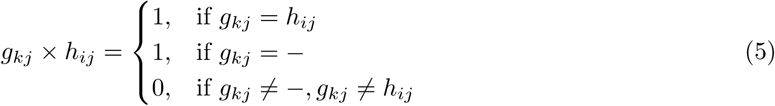

The fitness of a haplotype *h_i_* is then given as

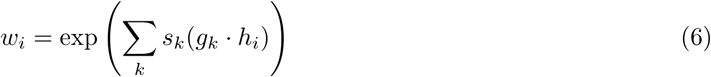

where the sum is calculated over all fitness parameters *k*. To give an example, a single-locus fitness parameter would have a single element of *g_k_* that was either A, C, G, or T. Supposing this element to be at position *j*, it would convey the fitness advantage *s_k_* to all haplotypes with the given nucleotide at position *j* in the genome.

#### Selection in a transmission event

Selection is incorporated into the transmission event from donor to recipient by representing this event as a biased sampling process. As we are not considering data here, noise is not an issue. We therefore model the population ***q**^F^* as arising via a multinomial sampling process of depth *N^T^* from a set of genotypes with frequencies *S^T^*(***q**^B^*), where *S^T^* represents the role of selection in the transmission event. We write

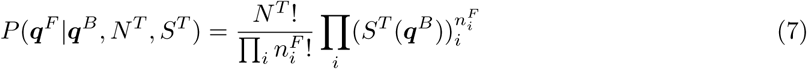

where

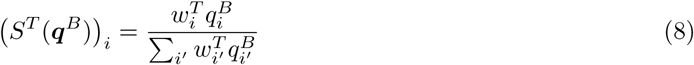

defines a distorted population based on the haplotype fitnesses 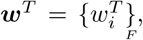 representing the relative propensity of each haplotype *h_i_*, for transmission. We note here that 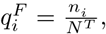 where ***n**^F^* is the composition of haplotypes in the founder population.

#### Selection during within-host growth

From the founding of an infection in the recipient, the viral population grows to the point at which data is collected for sequencing, under the influence of both genetic drift and selection. Selection for within-host growth is modelled by the function *S^G^*, identical in form to *S^T^*. We note that neglect of this term could distort the inferred value of *S^T^*; given only data collected before and after transmission the two terms cannot be separated. However, where samples have been collected at distinct times from one or multiple hosts, it is possible to make an independent estimate of *S^G^* [51], such that the two forms of selection can be discriminated. We here incorporate within-host selection into our derivation; the absence of such selection is then represented as a special case of our model.

Concerning drift, we note that the number of viruses in a host grows rapidly, with experiments suggesting that a single infected cell can produce between 10^3^ and 10^4^ viruses [73]. Each strand of RNA forming a new virus undergoes at least two rounds of replication within the cell; this replication has elsewhere been considered as a branching process with a mean 100-fold increase in the population size at each step [74]. In a population of variable size, the effective population size can be written as

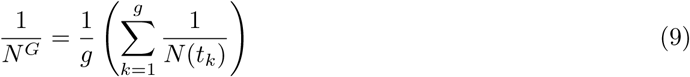

where *N*(*t_k_*) is the population size after k generations [75]. Given the rapid growth in population size we approximate the sum to consider only the first generation, modelling drift as a single multinomial draw with depth *N^G^* = 100*N^T^*.

#### Approximation of the likelihood function

We now turn to calculating the likelihood function of Equation 2. On account of the discrete nature of the multinomial distribution, the integrals present in this equation may be written as sums over all possible outcomes of the multinomial sampling processes represented by the different potential values of ***q**^F^* and ***q**^A^*. However, in realistic cases, where there might be multiple haplotypes present, the number of possible outcomes grows combinatorially with *N^T^*, making this calculation intractable. Instead we consider a continuous approximation in which the random variables of the model (Figure 2A) are represented by multivariate normal distributions, each defined by a mean and covariance matrix. By ignoring higher order moments, we may then calculate the individual components of the system (Equation 2) by appealing to a moments based approach for the evaluation of integrals arising from marginalisation over unknown variables. This step follows multiple previous approaches to time-resolved data, in which moments-based approximations have been used to simplify the propagation of evolutionary models [35,76–78].

The haplotype frequency vector ***q**^B^* is unknown and must be determined from the available data. We denote the mean of the distribution of ***q**^B^* as ***μ**^B^* and its covariance matrix by Σ^*B*^. Given a sampling depth *N^B^* and a dispersion parameter *C*, we describe ***x**^B^* as a distribution with mean and variance derived from the Dirichlet multinomial [79]:

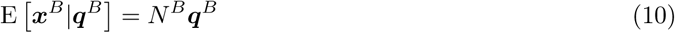

and

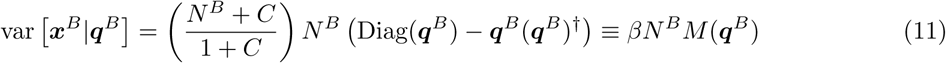

where 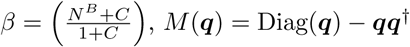 and † indicates the transpose function.

The founder population ***q**^F^* is sampled from ***q**^B^*. Its mean is given by the expression

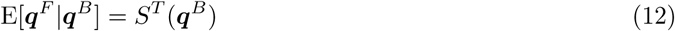

and its variance by

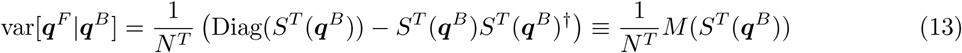

arising from a multinomial sample of depth *N^T^* and the selectively shifted frequencies *S^T^* (***q**^B^*).

Similarly, the within-host growth process may be represented by a distribution with mean E[***q**^A^*|***q**^F^*] = ***q**^F^* and variance var 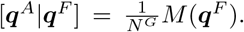 As for the pre-transmission case, a Dirichlet multinomial likelihood with sampling depth *N^A^*, selectively shifted frequencies *S^G^*(***q**^A^*) and dispersion parameter *C* may be used to model the sequencing of the population post-transmission. The resulting distribution can be approximated as a multivariate normal with mean

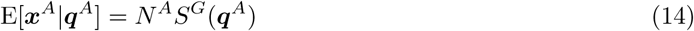

and variance

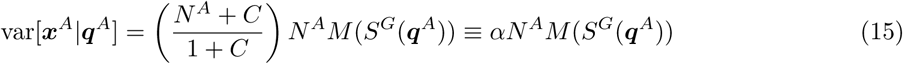

where 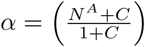 is defined for notational convenience.

Having established the above distributions, we are now equipped to carry out the relevant marginalisations (Equation 2) using the law of total expectation and the law of total variance. Starting with the pre-transmission compound distribution, the marginalisation over ***q**^B^* yields a mean of

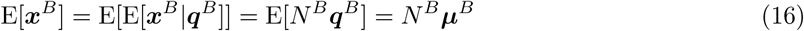

and a variance of

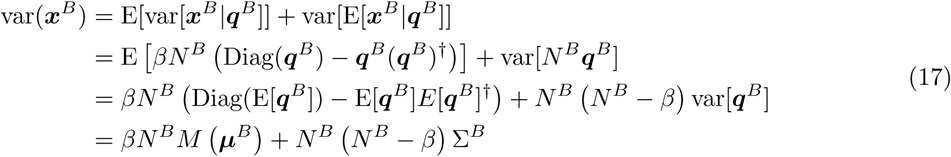

These expressions characterise the ***x**^B^* component of the likelihood from Eq. 2 in terms of a normal distribution. We identify values of ***μ**^B^* and Σ^*B*^ maximising this likelihood. As the covariance matrix Σ^*B*^ may contain a large number of elements, we make the approximation that its off-diagonal elements are zero.

Moving on to the post-transmission process, the marginalisation over ***q**^B^* results in a mean of

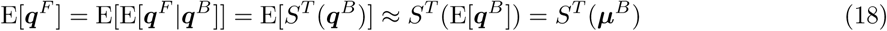

where in the penultimate step we used the first-order second-moment approximation to a vector function acting on a random variable. The law of total variance yields

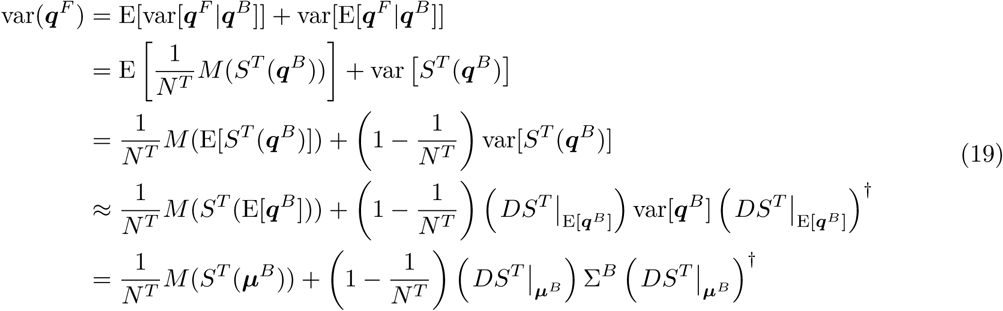

Note that 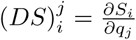 is the Jacobian matrix arising from the first-order second-moment approximation.

Marginalisation over ***q**^F^* yields a mean of

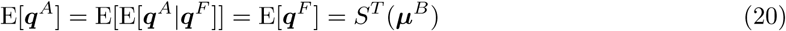

and variance

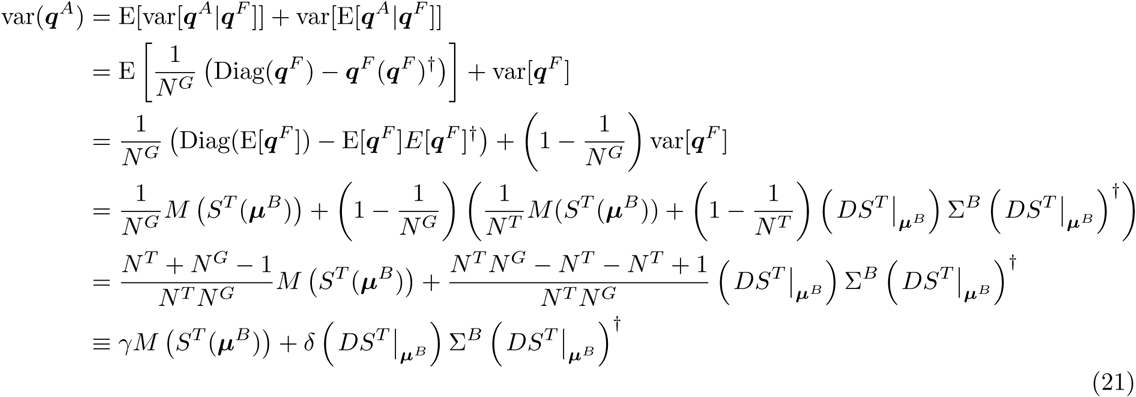

where in the last step we defined 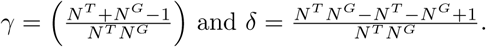

Treating the integral over ***q**^A^* in a similar manner, we obtain by the law of total expectation

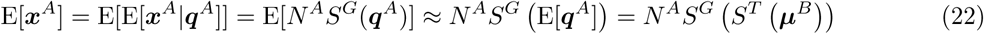

Analogously, the law of total variance yields

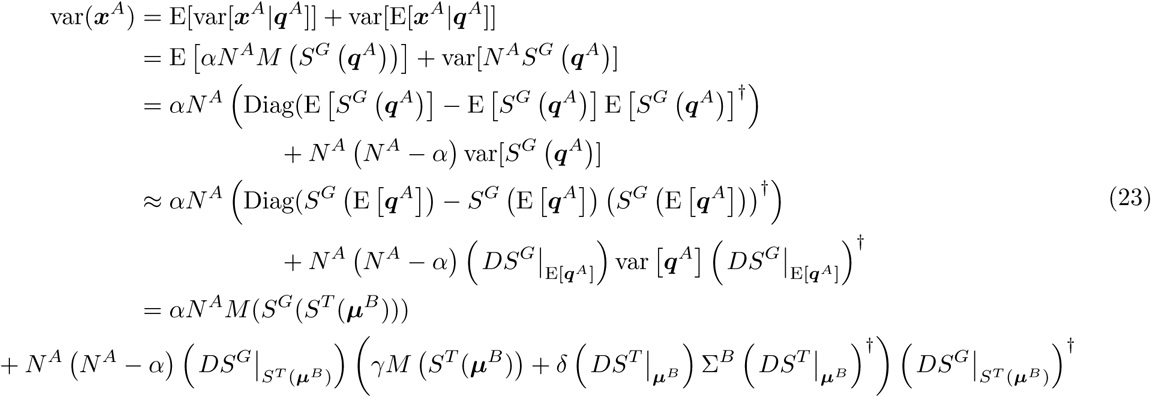

The above expressions represent mean and covariance matrices of multivariate normal distributions resulting from the evaluation of marginalisations in Equation 2. As such, the components of Equation 2 may be represented in a tractable form as the probability density functions of two multivariate normal distributions; The ***x**^B^* component has mean and covariance matrix as specified in Equations 10 and 11, whilst the ***x**^A^* component has mean and covariance matrix as given in Equations 22 and 23. Taken as a whole, this defines a likelihood for the transmission event given the data. As such, given an independent estimate of *S^G^*, and our estimated values for *μ^B^* and Σ^*B*^, the maximum likelihood values of *N^T^* and *S^T^* may be inferred.

### Reversion to a discrete likelihood function

Given a mean and covariance matrix for the likelihood function, we can approximate the likelihood by the probability density function of a multivariate normal distribution. However, where the variance of this distribution is very small in one dimension, as can occur under an inference of very strong selection, the density function evaluated at a point can become arbitrarily large. For this reason a Gaussian cubature approach was used to calculate the integral of the final likelihood over the unit cube described by each observation ***x***, once optimisation had been completed. Approximate numerical integrals were calculated using the software package cubature [80].

### Extension to partial haplotype data

In the calculations above we made the implicit assumption that the observations ***x**^B^* and ***x**^A^* consist of sets of complete viral haplotypes ***h**_i_*. However, short-read sequencing technologies generally result in sets of individual reads which only cover a subset of the genetic loci of interest; we refer to these reads as partial haplotypes. In this framework the data represents direct observations of partial haplotypes in the set 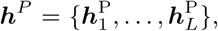 where each of the sets 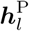 is a vector of haplotypes spanning a common subset of the loci spanned by the full haplotypes in ***h***. Population-wide observations of these partial haplotypes are then defined by 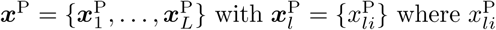 is the number of reads with haplotype 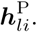 As a result, the total number of observations must now be defined on the basis of each set of partial haplotypes, e.g. 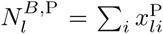 is the total number observations of partial haplotypes in the set *l*. As each set of partial haplotype observations is independent of the others, we may reconstruct Equation 2 in the following terms:

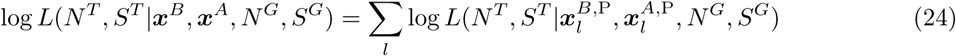

Within this construction, bottleneck sizes and selection are conserved between partial haplotype sets, being evaluated at the full haplotype level. Each set of partial haplotype observations 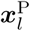 is considered as a sample drawn from a set of partial haplotypes with frequencies 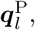 these frequencies being defined via a linear transformation of the full haplotype frequencies with matrix *T_l_*. For example, given the full haplotypes {AG, AT, CG, CT} and a set of partial haplotypes {A-, C-}, we have

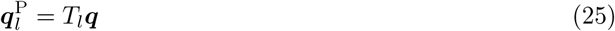

or more explicitly,

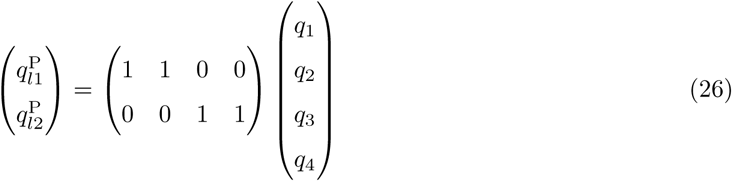

Thus, as described above, the calculation of transmission and within-host growth under selection can be performed at the level of full haplotypes, switching into partial haplotype space only to evaluate the likelihoods of the observations. Re-deriving the results of Equations 16 and 17 for short-read sequence data, we find that the compound distribution for the ***x**^B^* component has mean

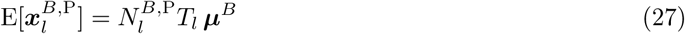

and variance

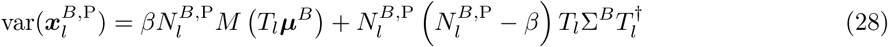

Similarly, for the ***x**^A^* component of the likelihood, we get a mean of

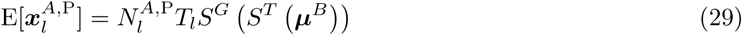

and variance

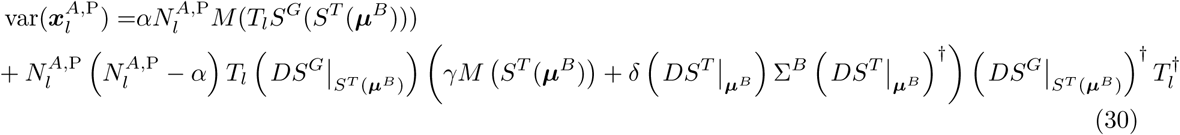

### Data from multiple genes

The mathematical framework outlined above utilises the haplotype information inherent to the data, and accounts for the effect of noise in the sequencing process (Figure 1B,C). However, in order to discriminate between changes in viral diversity arising from bottlenecking and selection (Figure 1A) it is necessary to consider data from different regions of the genome at which genetic diversity is nominally statistially independent. At high doses of influenza virus reassortment occurs rapidly, as has been observed both *in vitro* and in small animal infections [81,82]. In our analysis, distinct viral segments were therefore considered to be independent of one another in this manner, albeit sharing a common transmission bottleneck *N^T^*, each transmitted virus being assumed to contain one of each viral segment. As such the likelihood in Equation 24 becomes

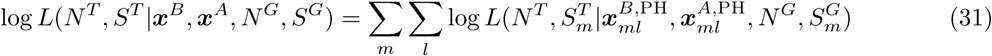

where the subscript *m* denotes information particular to a specific genomic region.

### Data from multiple replicates

Replicate data are highly valuable for evolutionary inference [83,84]. Within our calculation they provide an additional level of abstraction to the inference process. Under this framework we assumed that replicates share a common fitness landscape, *S^T^*, whilst exhibiting individual bottleneck values. As a result, the likelihood from Equation 31 becomes

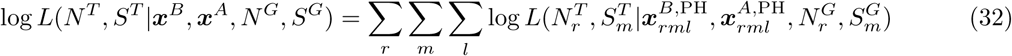

where the subscript *r* denotes information particular to a specific replicate.

### Application to data

Our method was applied to both simulated sequence data, and data from an evolutionary experiment conducted in ferrets [24].

#### Generation of simulated data

Simulated data were generated in order to nominally reflect data from an influenza transmission event. As such, a single transmission event was modelled as the transmission of viruses each with eight independent segments, the lengths of each segment being equal to the eight segments of the A/H1N1 influenza virus, with five randomly located polymorphic loci in each segment creating a total of 2^5^ potential full haplotypes. One fourth of these haplotypes were randomly chosen under the constraint that each of the five loci had to be polymorphic. Subsequently, full haplotype frequencies were generated at random, with the constraint of a minimum haplotype frequency of 5%.

Transmission was modelled as a multinomial draw of depth equal to the bottleneck size. Selection for transmission was incorporated as a shift in haplotype frequencies as described in Equation 8. Within-host growth was included as two 12-hour rounds of replication, each round being modelled as two successive drift processes, each resulting in a 100-fold increase in population size. Within-host selection was modelled in a manner similar to that of selection for transmission.

Partial haplotype observations were generated on the basis of short-read data simulated for each gene. Short-reads were modelled as randomly placed gapped reads with mean read and gap lengths derived from an example influenza dataset [23] (mean read length = 119.68, SD read length = 136.88, mean gap length = 61.96, SD gap length = 104.48, total read depth = 102825); these estimates are conservative relative to what can be achieved with the best contemporary sequencing technologies. Read depths were calculated for all possible sets of partial haplotypes by assigning individual reads to sets according to the loci they cover. Finally, partial haplotype observations were modelled as Dirichlet-multinomial draws employing a dispersion parameter *C* to account for noise.

Replicate experiments were generated by considering replicate observations of transmission events with consistent viral populations; that is, for which the variant alleles were consistent between replicate transmission events.

#### Experimental sequence data

Data were analysed from an evolutionary experiment in the transmission of a 1918-like influenza virus between ferrets [24]. The specific data examined here describes two sets of viral transmissions. In the first, denoted HA190D220D, a viral population was given to three ferrets, transmission to a recipient host being observed in one of three cases, giving time-resolved sequence data from four ferrets. In the second, denoted Mut, a viral population arising from the first experiment was given to three ferrets, transmission to two recipient hosts being observed, giving data from five ferrets.

#### Processing of sequence data

Genome sequence data was processed using the SAMFIRE software package, according to default settings [85], calling variant alleles that existed at a frequency of at least 1% at some point during the observed infections. For the calculation of a within-host fitness landscape, the effective depth of sequencing was estimated in a conservative manner, filtering out variants which changed in frequency by more than 5% per day before using the frequencies of remaining variants from different time-points within the same host to estimate the parameter *C*. Following the approach of previous calculations [51,69], potentially non-neutral variants were identified as those for which a model of frequency change under selection outperformed a neutral model by more than 10 units according to the Bayesian Information Criterion (BIC) [56]. Variants reaching a frequency of at least 5% in at least one sample were then identified before calling multi-locus variant observations from the data; data from all time-points were used in this inference. The 5% cutoff was chosen to reduce computational costs for this part of the calculation while still reconstructing the core aspects of the within-host fitness landscape.

For the inference of transmission, a revised approach to estimating the effective depth of sequencing was taken, noting our result that estimates which overestimate noise may lead to errors in the inferred bottleneck size. Here, in common with previous calculations, we initially identified a conservative value of *C* from within-host data using the default settings in SAMFIRE. Next, variant frequencies were evaluated, identifying potentially non-neutral changes in frequency using a single-locus analysis [51]. Finally, a more conservative estimate of *C* was calculated, using the set of trajectories which were identified as being consistent with a neutral model of frequency change.

As in the original analysis of the data [24], variants were identified from data collected from the final observation before transmission and the first point of observation after transmission; these data were used to construct multi-locus observations across variants which reached a frequency of at least 2% in at least one sample.

Subsequent processing was identical for simulated and experimental datasets: Partial haplotype observations were removed if A) the partial haplotype did not have at least 10 observations either before or after transmission, B) the partial haplotype exhibited a frequency of < 1% before transmission, C) the partial haplotype had no observations before transmission (variant assumed to have arisen de novo), D) the partial haplotype was the only partial haplotype in its set and had no observations post-transmission. Additionally, to avoid potential dataset errors from drastically influencing the inference outcome, partial haplotypes were removed if found to have a single post-transmission observation despite the presence of a large (≥ 50) overall sampling depth. Finally, removal of partial haplotype observations may result in individual loci becoming monomorphic (all partial haplotypes covering these loci exhibit the same alleles). In this case, relevant partial haplotype sets were removed with the reads being redistributed unto variant sets with fewer loci.

### Inference of parameters

#### Hierarchical selection model

In our model, the set of potential fitness parameters is large. To simplify the calculation, parameters modelling three- or higher-locus epistatic effects were neglected, while parameters modelling two-locus epistasis were only considered for addition to a model which already contained single-locus fitness parameters for each of the two loci. In both the inferences of within-host selection and of transmissibility, a null assumption of neutrality was used as the starting point for an inference model, exploring successively more complex models of selection until an optimal model, defined according to a model selection process, was identified.

#### Inference of within-host selection

For the experimental dataset an inference of within-host selection was conducted according to a method previously described in earlier publications [51,69]. Under the assumption of rapid reassortment in the system [81] different segments of the virus were treated independently. Our inference of selection aimed to characterise fitness so as to estimate *S^G^* for an inference of transmission; the HA190D225D and Mut datasets were considered independently, with data from all animals in each set being combined to infer within-host selection.

#### Replicate calculations of transmission parameters

Both our within-host and transmission calculations are performed in a model space of potential multilocus haplotypes, identified by the SAMFIRE code [85]. In the first step of the transmission model, we calculate an estimate for the population ***q**^B^* given the data *x*^*B*^. Where there are greater numbers of potential haplotypes and short reads span smaller numbers of loci there is an increasing potential for the data to not fully specify the initial vector ***q**^B^*. For this reason statistical replicate calculations were run in each case, using different reconstructions of ***q**^B^* in each case; the median inferred parameters across replicates are presented above. To improve the speed of the inference, haplotypes in ***q**^B^* with inferred frequencies of less than 10^−10^ were removed from the calculation; subsequent to this, haplotypes were removed in increasing size of inferred frequency until no more than 100 haplotypes remained in ***q**^B^* at non-zero frequencies. We note that our inference of ***q**^B^* depends upon the initial identification of a plausible set of underlying viral haplotypes using SAMFIRE. Where this plausible set is large, as might occur where very short reads describe a large number of polymorphic loci, this inference becomes computationally challenging.

#### Model selection

Model selection was performed using the Bayesian Information Criterion:

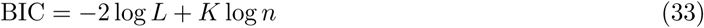

where *L* is the maximum likelihood obtained for a model, *K* is the number of parameters in the fitness model, and *n* is the number of data points. A range of potential fitness models were explored, the optimal model being identified as that for which the addition of any single fitness parameter failed to bring a significant improvement in BIC.

#### Adaptive BIC

Noting previous discussion of the complexity of using BIC in biological modelling [86], we here adopted a machine-learning approach to the interpretation of BIC statistics. Classically, a difference of 10 units of BIC has been held to represent strong evidence in favour of the additional parameter [56]. Consistent with previous approaches this heuristic was used in the inference of within-host selection; in this case the final model parameters make only small refinements to the inferred fitness landscape [51]. In the inference of transmission, errors in model selection have more severe consequences for the inferred bottleneck size and selection model. Using a fixed difference of 10 BIC units for model selection resulted in an overestimation of the extent of selection with a high false positive rate (Supplementary Figure S12). As such, we generated and analysed simulated data to identify the optimal interpretation of BIC differences. Given a real dataset for analysis, simulated data was generated describing systems with equivalent numbers of gene segments and polymorphic loci to the real dataset, being observed with an equal number of reads spanning each set of loci, and with reads containing an amount of information specified by the parameter *C* inferred for the real dataset.

Next, inferences were conducted on data describing neutral transmission events with bottlenecks in the range [5,100]. As shown in Figure 3, the ability to infer a correct neutral bottleneck is impaired by noise for transmission events involving a large number of viruses; linear regression was used to obtain a simple function describing the ratio between the median inferred and true bottleneck sizes under neutrality (Supplementary Figure S13A); this parameterises our expectation of the ‘correct’ inferred bottleneck size for any given real bottleneck, once noise is accounted for.

Secondly, using this baseline to set our expectations, a parameterisation was carried out to find a BIC penalty function that gave the largest accuracy in bottleneck inference. To this end, three datasets were considered; a neutral dataset and two datasets with single selection coefficients of *s* = {1, 2} respectively. BIC penalty values in the range [10, 200] were examined, with smaller BIC penalty values leading to inferences with a larger number of selection coefficients and vice versa. For each BIC penalty value, the difference between the bottleneck inference of the optimal model (under BIC) and the baseline expectation was summed for the three datasets to give a statistic describing the accuracy of the inferred bottlenecks, this statistic being expressed as a function of the real transmission bottleneck *N^T^* and the BIC penalty (Supplementary Figure S13B). Finally, linear and decay exponential models were fitted to this data via regression, selecting the BIC penalty model which minimised the error in the inferred bottlenecks from the simulation data. We note that our penalty is a function of the inferred population bottleneck, higher penalties being inferred for tight bottlenecks and lower penalties being inferred for looser bottlenecks.

Thirdly, the inferred data was reinterpreted to derive a BIC penalty optimal for the inference of selection. We note that, even with a BIC penalty function optimised for bottleneck inference, there may still remain cases where, through the stochastic process of transmission, the genetic composition of the population changes in a manner consistent with the action of selection, granting a false positive inference. A second BIC penalty was learned as above, this time maximising the accuracy of the inference or non-inference of selection parameters, defined as

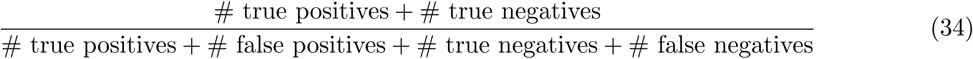

This conservative BIC penalty function typically led to an underestimate for the inferred bottleneck; the two BIC penalty functions were used in concert to estimate *N^T^* and *S^T^* in separate calculations.

As noted elsewhere, where a genomic variant fixes between two observations, this change in frequency can be explained by the fitting of an arbitrarily large selection coefficient; no upper bound on selection can be established [87]. Within our framework, if this is not accounted for, extremely strong selection may be falsely inferred to explain the loss of variants during a transmission bottleneck. To guard against this, models of transmission in which the inferred magnitude of selection was outside of the range (−10,10) were excluded from consideration. In the within-host analysis methods, haplotype fitness are not constrained; here, to avoid errors of machine precision, the magnitudes of extreme fitness inferences were reduced to be within the range (−10,10). For the same reason, elements of the mean and covariance matrix of ***q**^B^* were constrained to be greater in magnitude than 10^−11^.

## Acknowledgments

CJRI is supported by a Sir Henry Dale Fellowship jointly funded by the Wellcome Trust and the Royal Society (Grant Number 101239/Z/ 13/Z). CKL is supported by a PhD Studentship funded by the Wellcome Trust (Grant Number 105365/Z/14/Z).

## Supplementary Material

**Figure S1.**
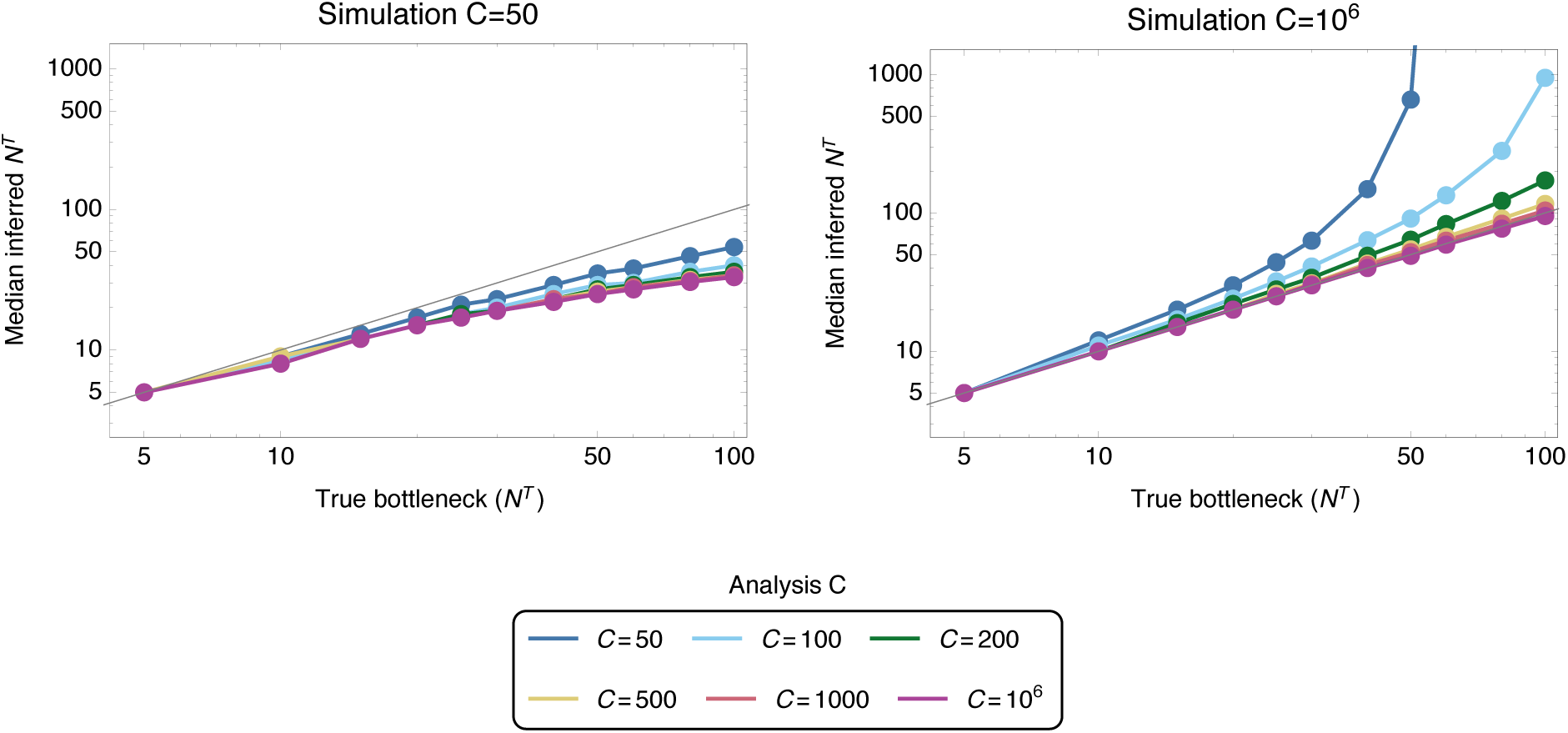
Bottleneck inference under a neutral model applied to neutral data with simulation dispersion parameters of *C* = {50,10^6^}. Inference was performed using a range of dispersion parameters, *C* = {50,100, 200, 500,1000,10^6^}.

**Table S1.** Inferred fitness coefficients for the within-host evolution of the virus within each experiment. Parameters were inferred across all index and contact ferrets within each experiment and are reported to a single decimal place. Only polymorphisms at which within-host selection was identified are listed. The parameter χ denotes an epistatic interaction between variant alleles. We note that our method infers the approximate shape of a fitness landscape based upon a reconstruction of whole viral segments; individual selection coefficients may be subject to variance between similar fitness landscapes.

| Segment | Variant | Mut | HA190D220D |
| --- | --- | --- | --- |
| PB2 | A1199G | -0.9 |  |
| PB2 | T1537C |  | 0.5 |
| PB2 | G2193C |  | -0.3 |
| PB1 | C65T |  | -0.4 |
| PB1 | C90A | -0.3 | -0.3 |
| PB1 | C835A |  | 0.3 |
| PB1 | G982T |  | 0.4 |
| PB1 | T1151G |  | 0.3 |
| PB1 | G2250T | -1 |  |
| PB1 | $\chi_{90,982}$ | | 1 |
| PA | A781G | -0.2 |  |
| PA | G1500T |  | 0.5 |
| PA | C1651T | -0.7 |  |
| PA | G1880T | -0.8 |  |
| HA | G14T | -0.0 | 0.4 |
| HA | A400G | 0.2 | -0.3 |
| HA | A507C | 0.5 | 0.4 |
| HA | C550A | 0.3 |  |
| HA | T634C | 0.4 |  |
| HA | A649G |  | 0.3 |
| HA | A651C | 0.6 |  |
| HA | T653G | 0.5 |  |
| HA | G741A |  | -0.1 |
| HA | G747A | 0.2 |  |
| HA | A748G | 0.3 |  |
| HA | A868T | 0.2 | 0.3 |
| HA | T1036C |  | -2.1 |
| HA | G1263A |  | 0.5 |
| HA | C1762T | -0.7 |  |
| HA | $\chi_{14,400}$ | -0.3 | |
| HA | $\chi_{14,507}$ | 0.2 | |
| HA | $\chi_{400,507}$ | -36.5 | |
| HA | $\chi_{400,550}$ | -32.2 | |
| HA | $\chi_{400,1036}$ | | 2.2 |
| HA | $\chi_{868,1263}$ | | -0.8 |
| NP | G600A |  | 0.4 |
| NA | G440A |  | 0.4 |
| NA | G649A |  | 0.2 |
| NS | G289A |  | 0.4 |

**Figure S2.**
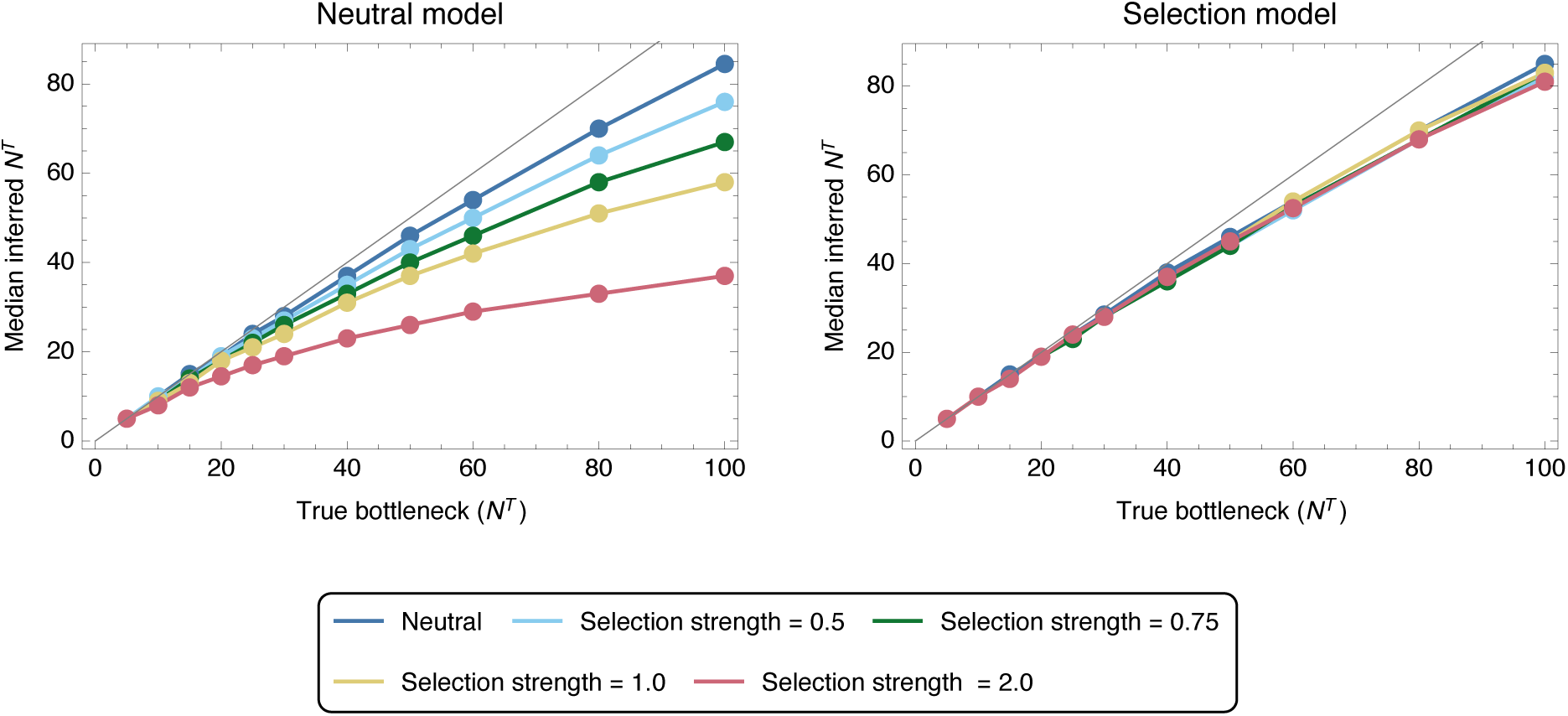
Median inferred bottleneck size from data simulating neutral transmission and transmission with a single locus under selection of magnitude *σ* ∈ {0,0.5, 0.75,1.0, 2.0}. Inferences were made using either a neutral model, in which the effect of selection was assumed to be zero, or a selection model, which allowed scenarios involving selection to be identified. Median inferences are shown from 100 simulations, each involving three replicate transmission events, for each datapoint.

**Figure S3.**
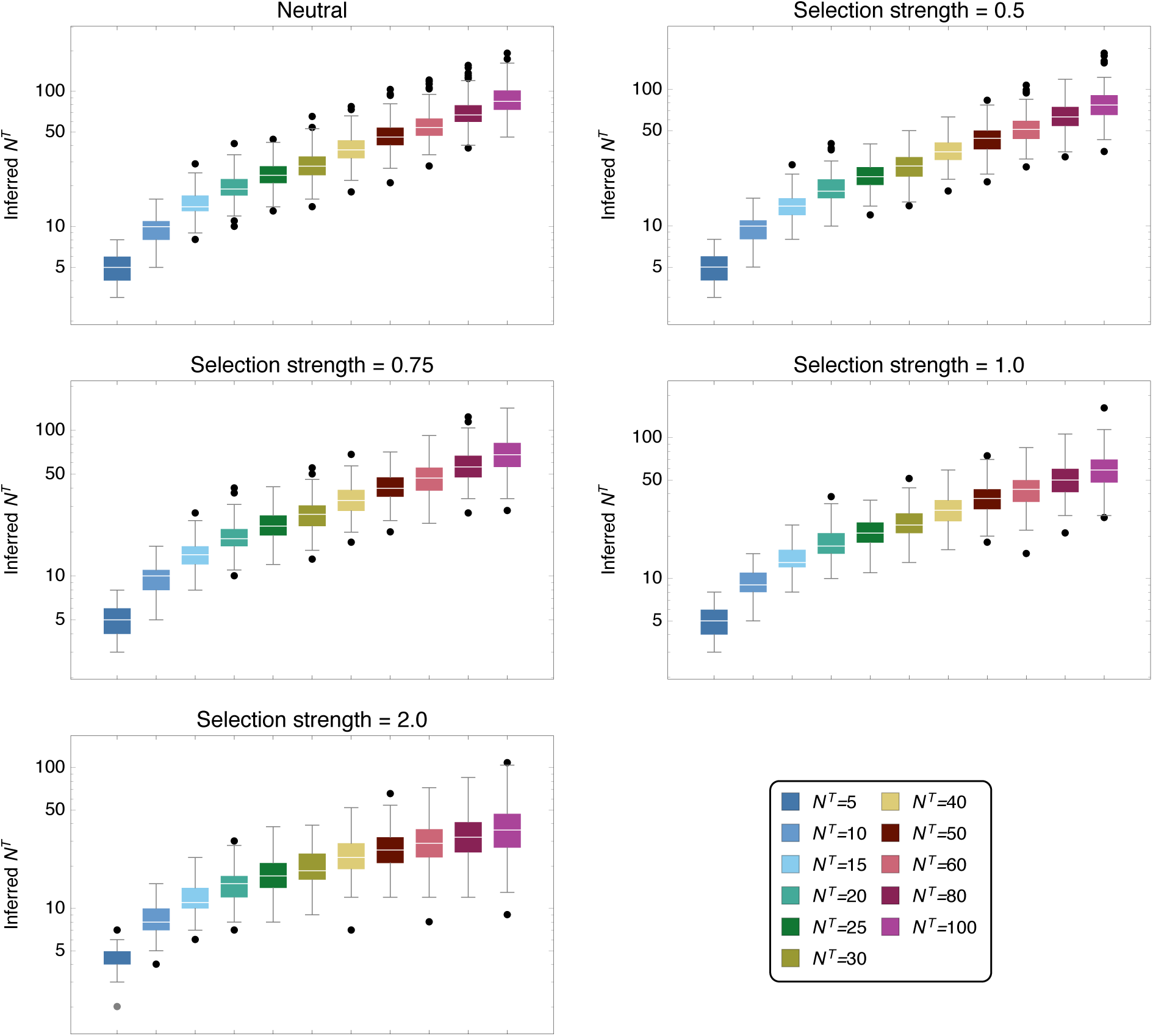
Inferred bottleneck sizes *N^T^* for a range of true bottleneck sizes. Results were generated by applying a neutral inference model to selected simulated data. Results are shown for 200 simulations at each bottleneck size.

**Figure S4.**
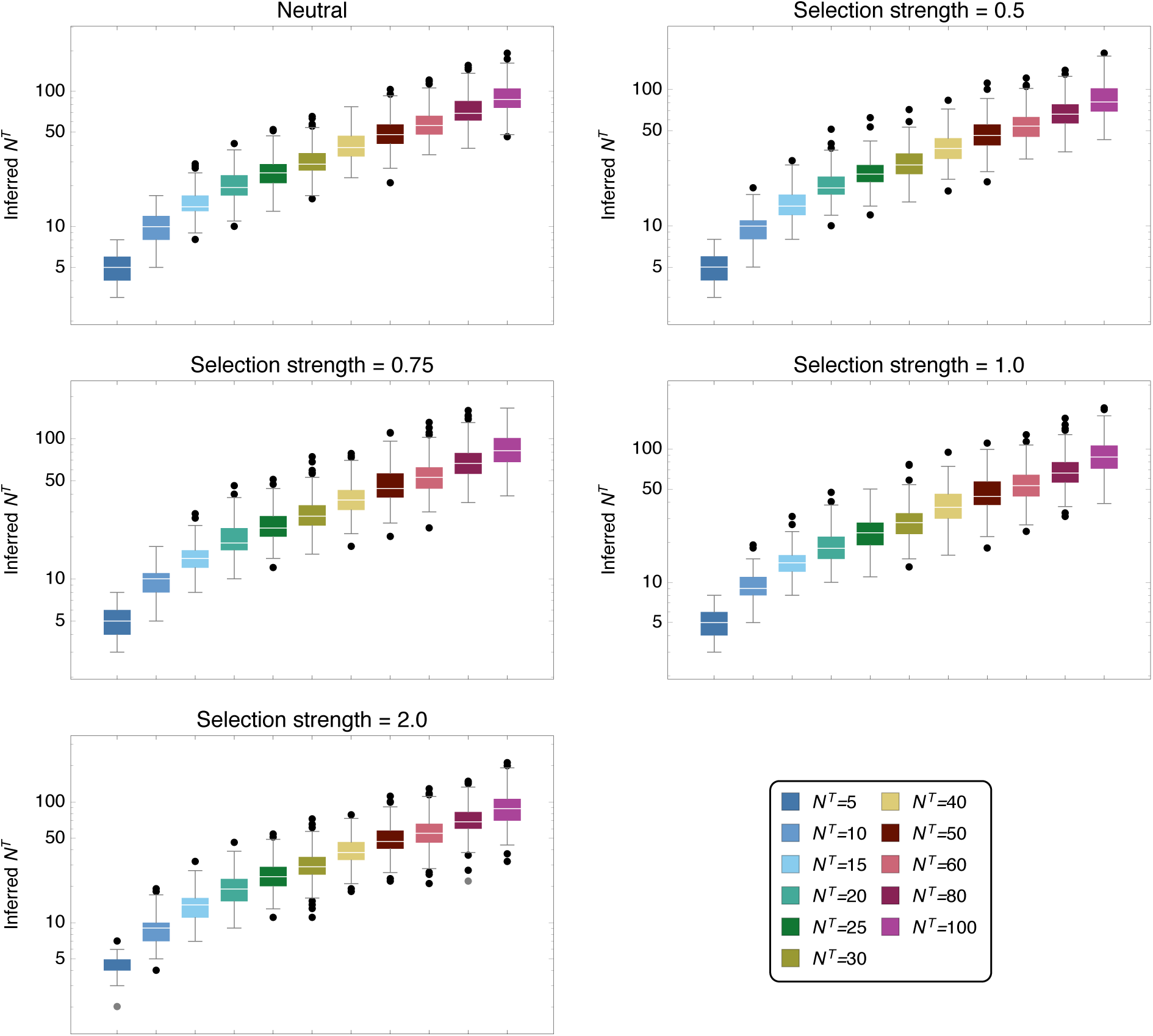
Inferred bottleneck sizes *N^T^* for a range of true bottleneck sizes. Results were generated by applying an inference model accounting for selection to selected simulated data. Results are shown for 200 simulations at each bottleneck size.

**Figure S5.**
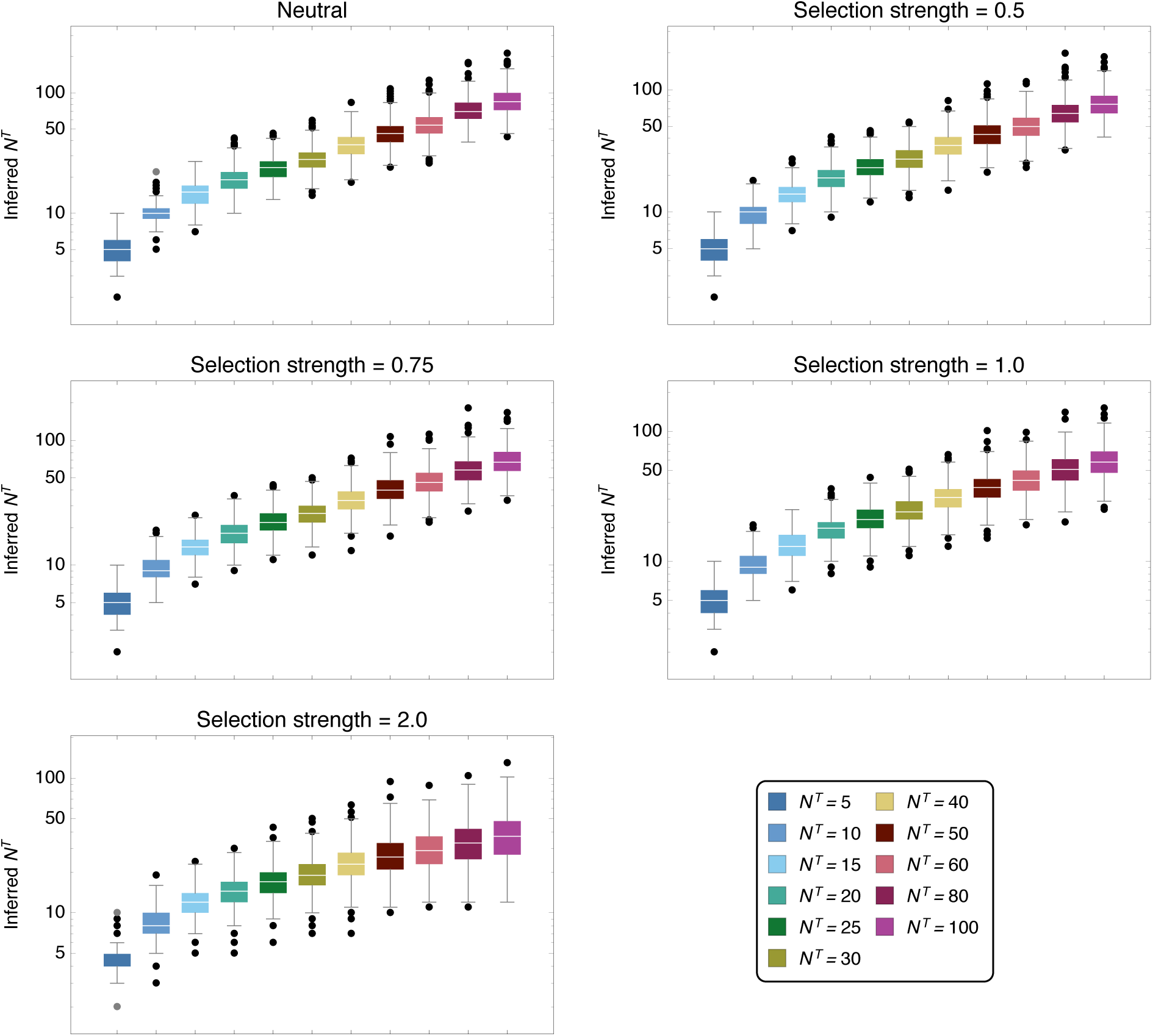
Inferred bottleneck sizes *N^T^* for a range of true bottleneck sizes. Results were generated by applying a neutral inference model to selected simulated data. Results are shown for 200 simulations at each bottleneck size, each simulation describing three replicate transmission events.

**Figure S6.**
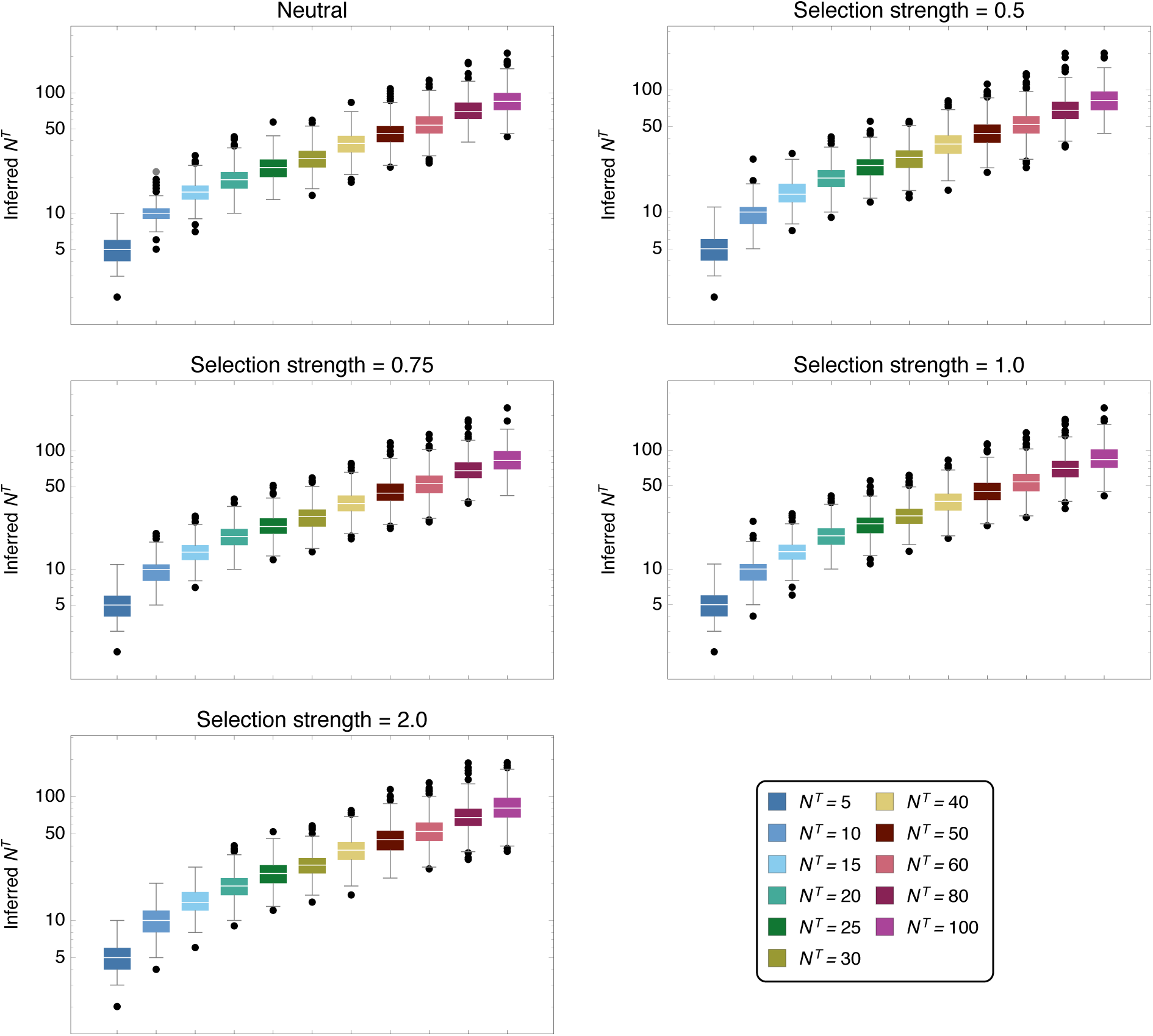
Inferred bottleneck sizes *N^T^* for a range of true bottleneck sizes. Results were generated by applying an inference model accounting for selection to selected simulated data. Results are shown for 200 simulations at each bottleneck size, each simulation describing three replicate transmission events.

**Figure S7.**
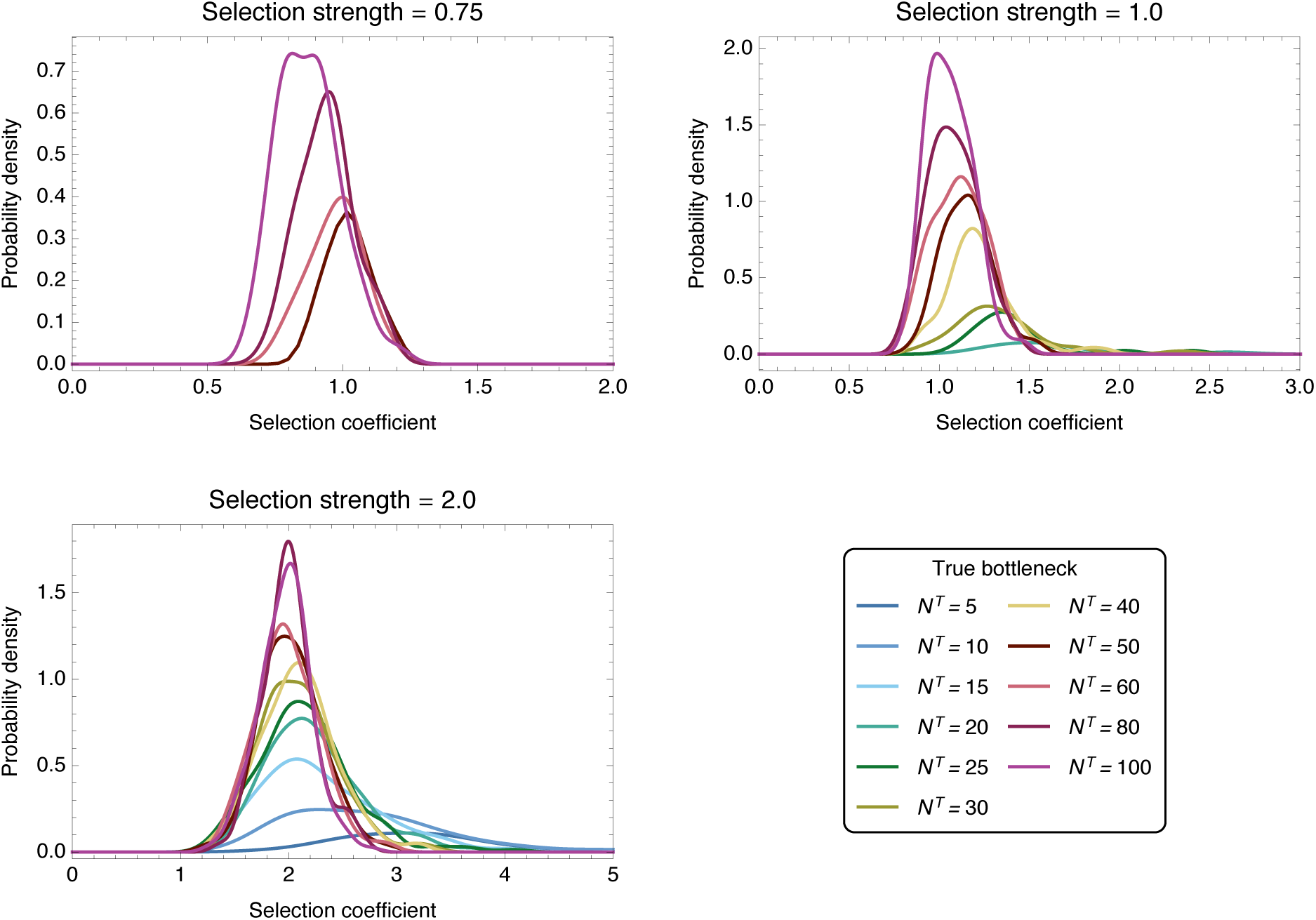
Probability distributions of inferred selection coefficients from 100 simulations each of three transmission events with selective pressures *σ* ∈ {0.75,1.0, 2.0}. Distributions were constructed for bottleneck values where the inference of selection resulted in a true positive rate for identifying selected variants of above 5 %. Smooth kernel distributions were computed as for Figure 7

**Figure S8.**
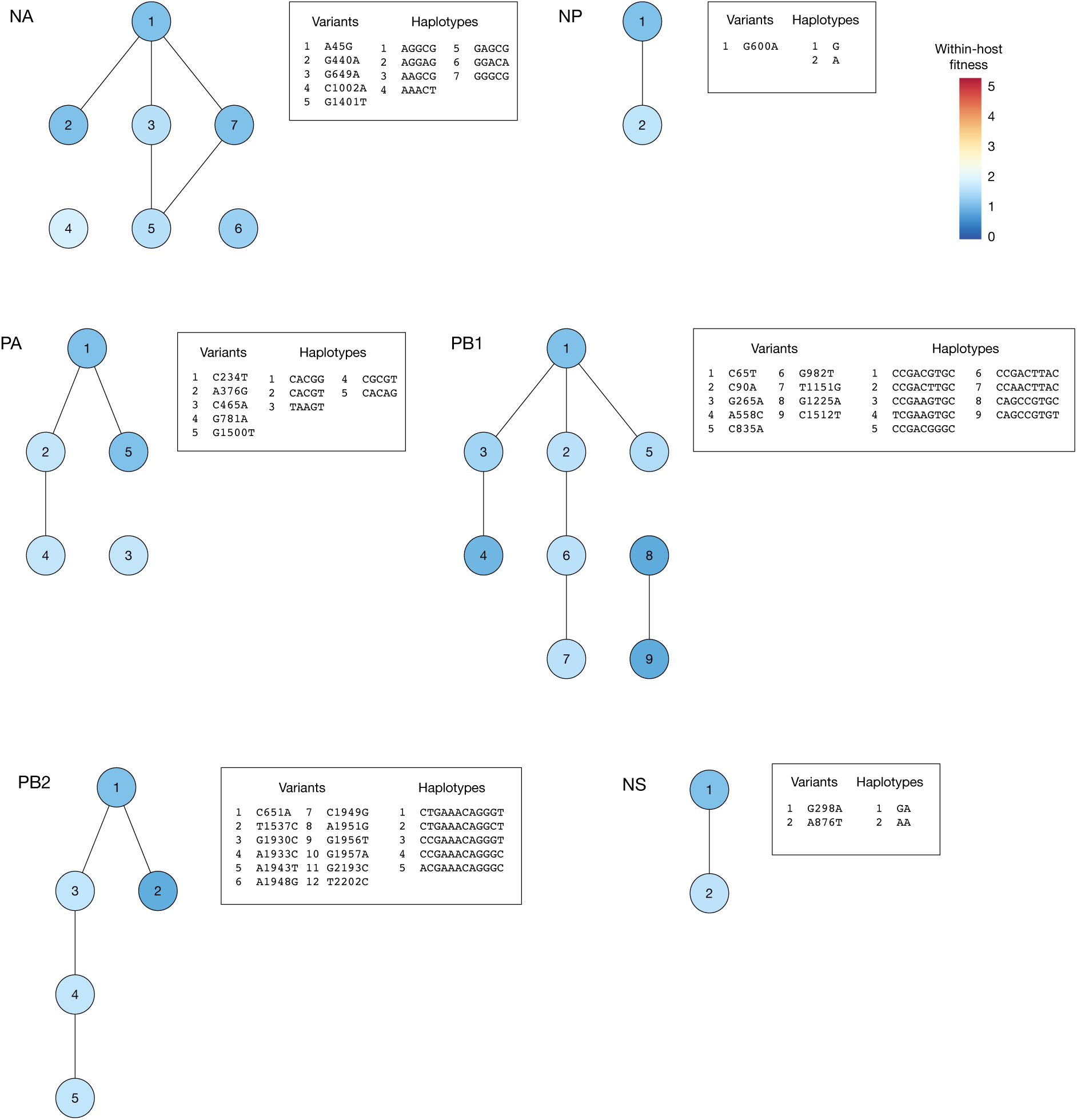
Inferred within-host fitness landscape for segments in the HA190D220D viral populations. Haplotypes for which the inferred frequency rose to a frequency of at least 1% in at least one animal are shown. Haplotypes which are separated by a single mutation are joined by lines.

**Figure S9.**
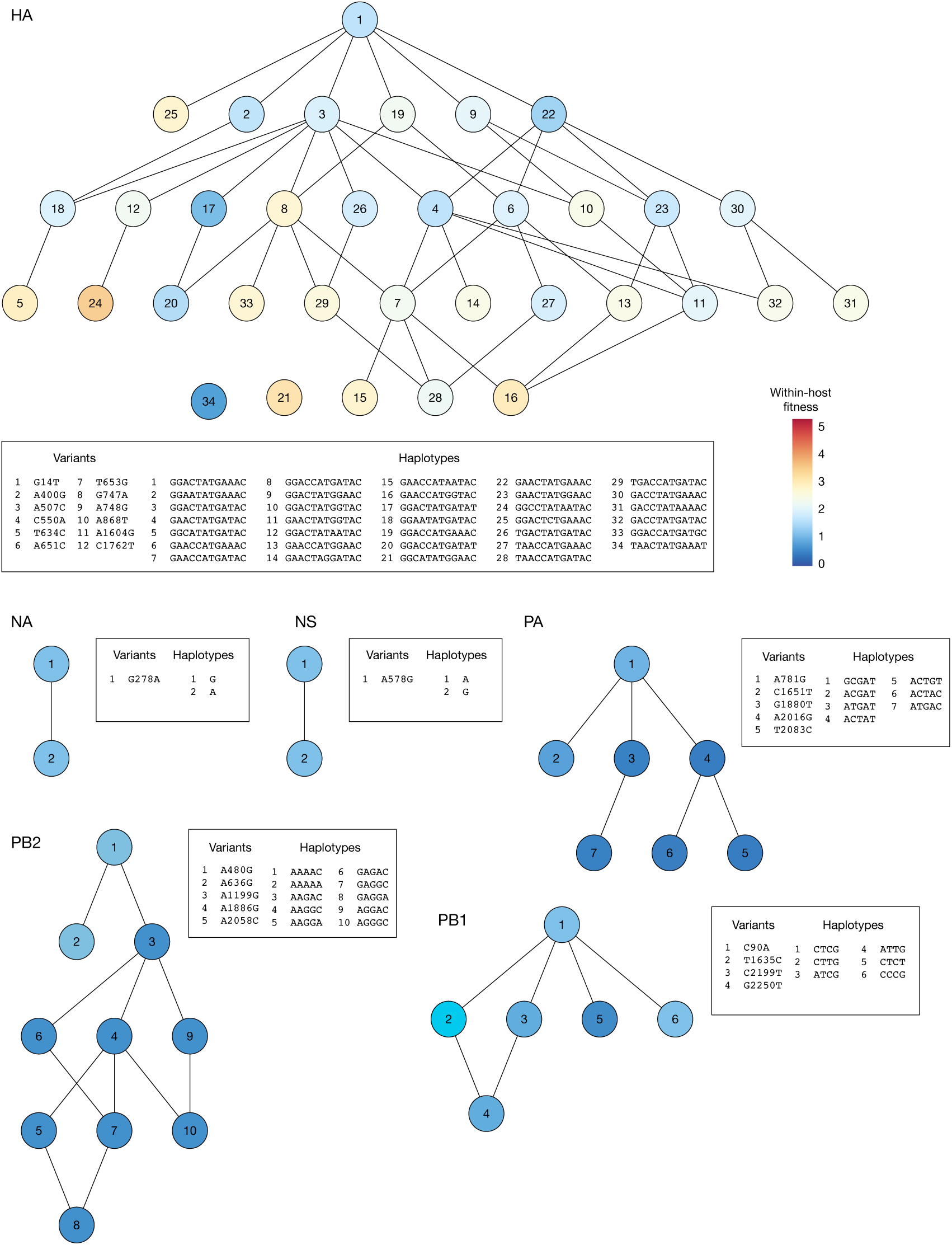
Inferred within-host fitness landscape for segments in the Mut viral populations. Haplotypes for which the inferred frequency rose to a frequency of at least 1% in at least one animal are shown. Haplotypes which are separated by a single mutation are joined by lines.

**Figure S10.**
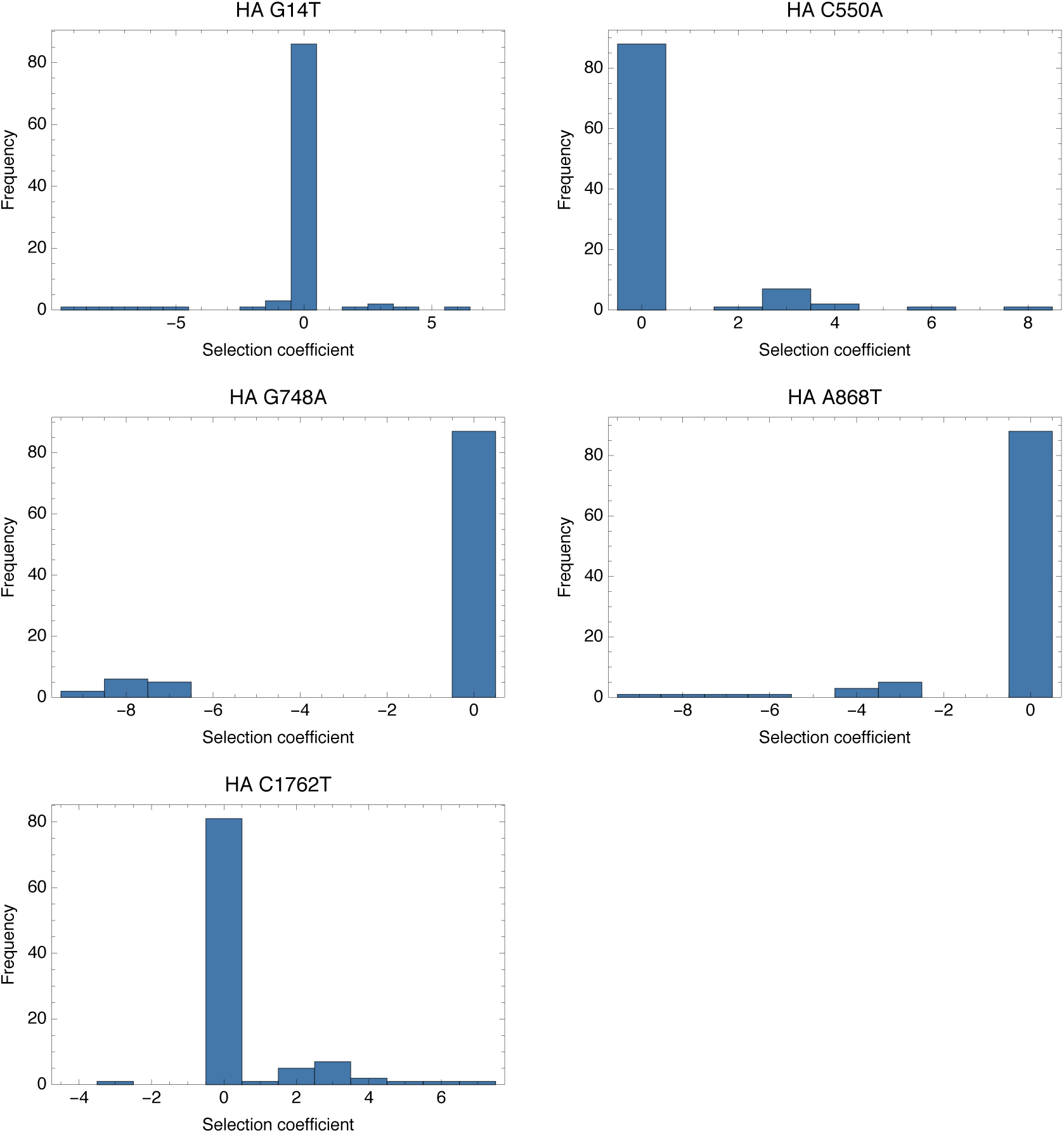
Histograms of selection inferences for the Mut transmission pairs from 100 random seeds using an allele frequency cut-off of 2%. A replicate inference method was employed such that a common fitness landscape was imposed. Selection inferences that resulted in at least 10% non-zero inferences are here reported by the nucleotide position of the variant site.

**Figure S11.**
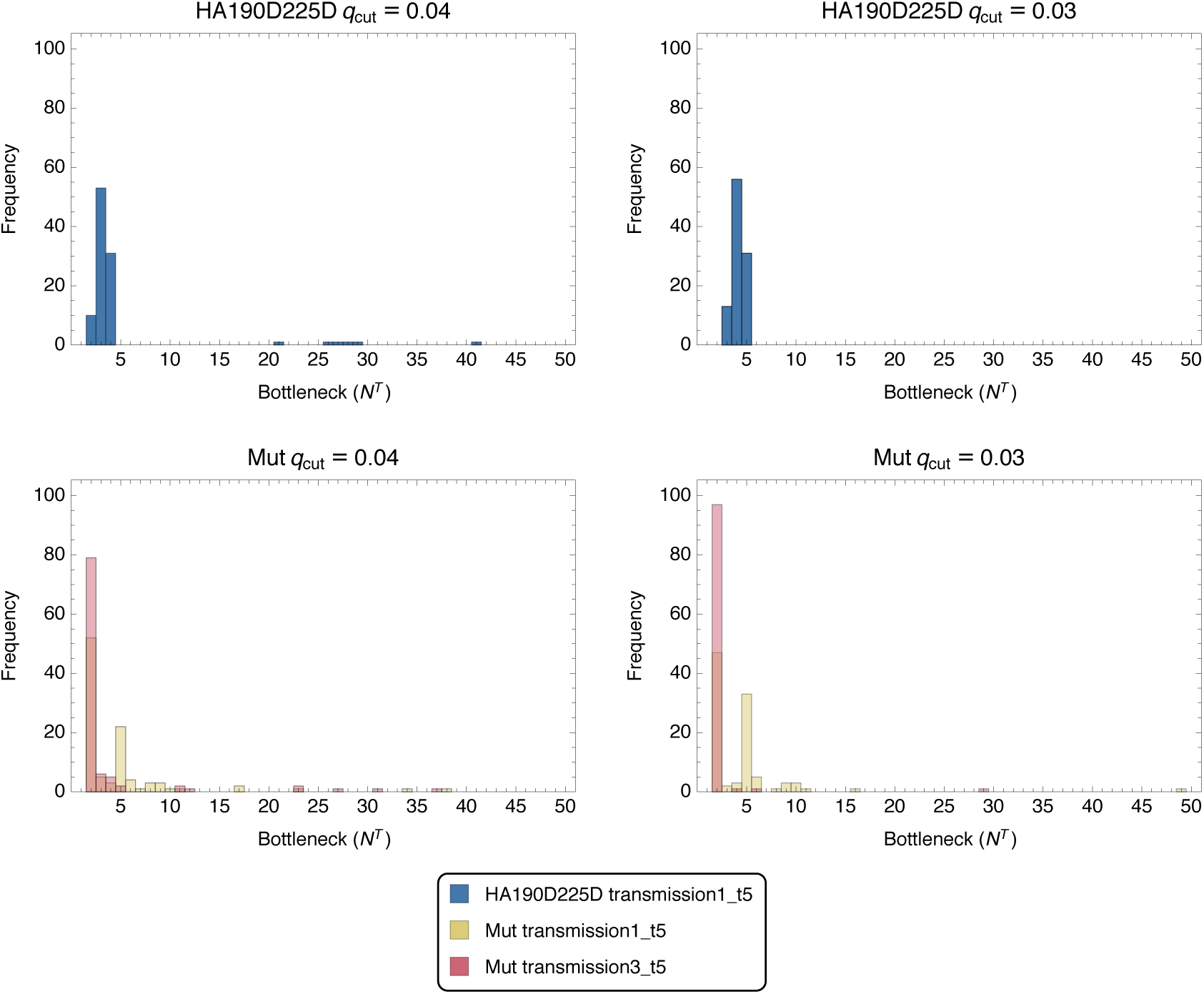
Histograms of bottleneck inferences for HA190D225D and Mut transmission pairs from 100 random seeds using allele frequency cut-offs of *q*_cut_ ∈ {0.03,0.04}. A replicate inference method was employed for the Mut transmission pairs such that a common fitness landscape was imposed. The Mut transmission pairs may take different bottleneck values and have been plotted as an overlapping histogram.

**Figure S12.**
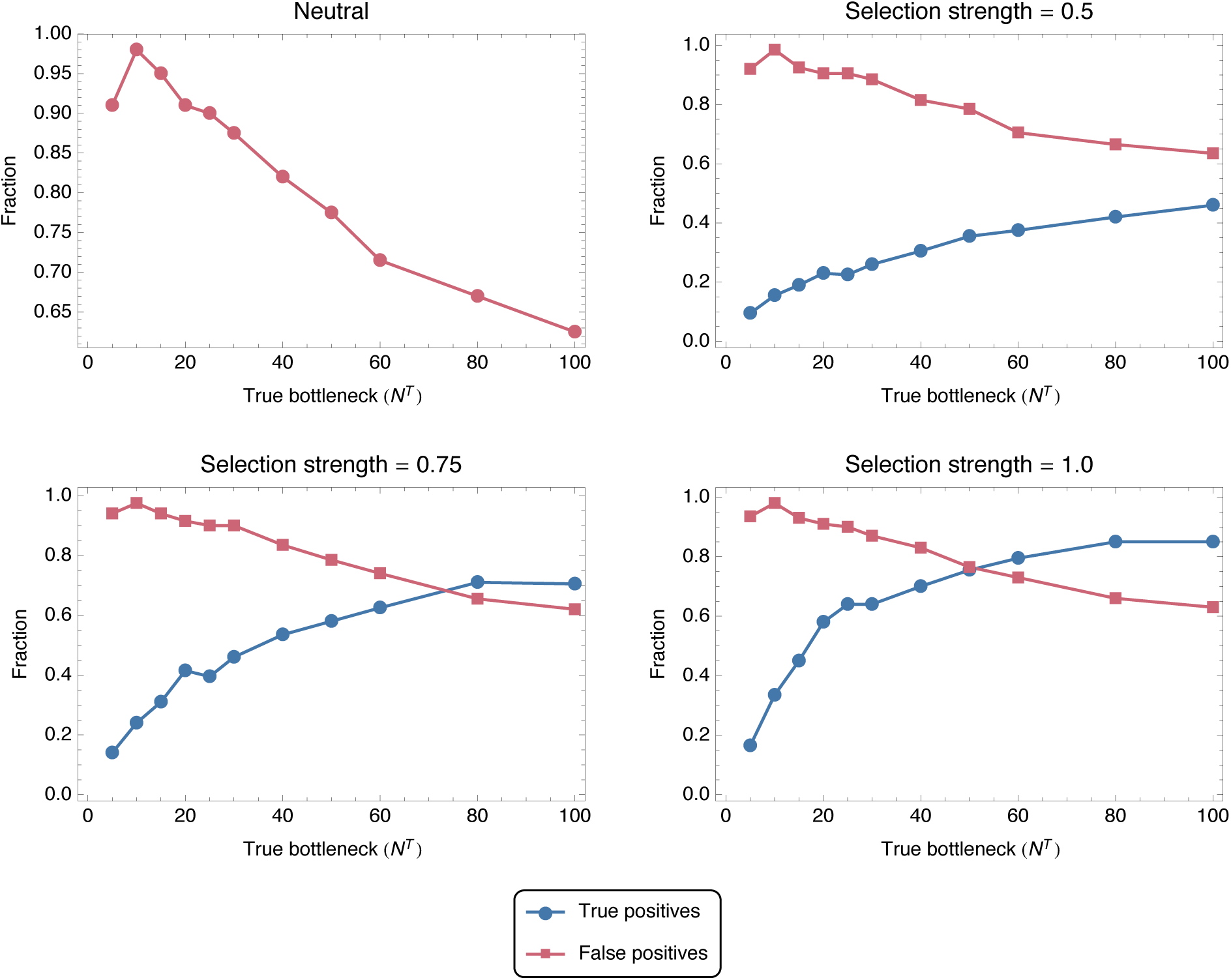
True and false positive rates of selection inference from 200 simulations of transmission events from single-replicate systems with selective pressures of *σ* ∈ {0,0.5, 0.75.1.0}. A fixed BIC difference of 10 units were employed in the model selection process, requiring a model with a single additional parameter to generate an improvement of at least 10 units to BIC to be accepted. While such a difference is accepted as showing strong evidence in favour of the more complex model, in our case it generated a high rate of false positive inferences of selection.

**Figure S13.**
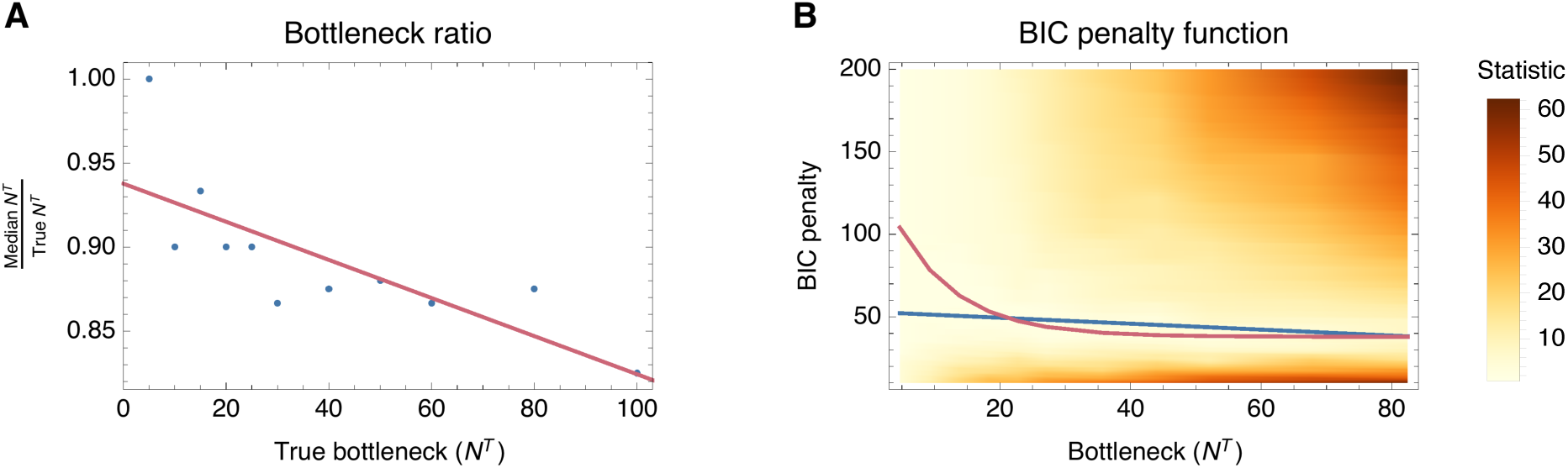
Determining BIC penalty function for bottleneck inference under simulated data. A) The ratio of the median inferred bottleneck to the true bottleneck is plotted against the true bottleneck size. As shown in Figure 3, as the bottleneck increases, our ability to infer it correctly decreases due to noise. In order to account for this phenomenon, a straight line is fitted to the data aiming to capture the general trend. B) Heat map of the bottleneck-specific statistic plotted against BIC penalty and bottleneck size. The plot was generated for three datasets with selection coefficients *s* = {0,1, 2} and a simple statistic based on bottleneck differences was employed. More specifically, the median bottleneck was computed across 200 seeds and the bottleneck-statistic was defined as the absolute value of the difference between the median inferred bottleneck and the true bottleneck multiplied by the baseline determined in A). By considering bottlenecks in the range [5,100] and BIC penalty values in the range [10, 200], a heat map was produced and linear and decay exponential regression were conducted seeking to minimise the sum of the statistic across the values of *N^T^* that were considered.

